# Thiolutin has complex effects in vivo but is a direct inhibitor of RNA Polymerase II in vitro

**DOI:** 10.1101/2021.05.05.442806

**Authors:** Chenxi Qiu, Payal Arora, Indranil Malik, Amber J. Laperuta, Emily M. Pavlovic, Scott Ugochukwu, Mandar Naik, Craig Kaplan

## Abstract

Thiolutin is a natural product transcription inhibitor with an unresolved mode of action. Thiolutin and the related dithiolopyrrolone holomycin chelate Zn^2+^ and previous studies have concluded that RNA Polymerase II (Pol II) inhibition *in vivo* is indirect. Here, we present chemicogenetic and biochemical approaches to investigate thiolutin’s mode of action in *Saccharomyces cerevisiae*. We identify mutants that alter sensitivity to thiolutin. We provide genetic evidence that thiolutin causes oxidation of thioredoxins *in vivo* and that thiolutin both induces oxidative stress and interacts functionally with multiple metals including Mn^2+^ and Cu^2+^, and not just Zn^2+^. Finally, we show direct inhibition of RNA polymerase II (Pol II) transcription initiation by thiolutin *in vitro* in support of classical studies that thiolutin can directly inhibit transcription *in vitro*. Inhibition requires both Mn^2+^ and appropriate reduction of thiolutin as excess DTT abrogates its effects. Pause prone, defective elongation can be observed *in vitro* if inhibition is bypassed. Thiolutin effects on Pol II occupancy *in vivo* are widespread but major effects are consistent with prior observations for Tor pathway inhibition and stress induction, suggesting that thiolutin use *in vivo* should be restricted to studies on its modes of action and not as an experimental tool.

## Introduction

Thiolutin is a historically used transcription inhibitor with potentially multiple modes of action. Thiolutin one of the microbially produced dithiolopyrrolone compounds that feature an intra-molecular and redox-sensitive disulfide bond (1,2). Holomycin is another notable compound in this class. These compounds are thought to be pro-drugs whose anti-microbial action requires reduction of the intramolecular disulfide. Thiolutin has been shown to inhibit bacterial and eukaryotic transcription *in vivo* and has been used to study mRNA stability in multiple species (3–6). However, the mode of action of thiolutin for transcription inhibition remains unclear and complicated (discussed below). In addition, thiolutin appears to affect multiple cellular pathways, including glucose metabolism (7), Tor signaling (8), Hog/MAPK pathway signaling (8), P body formation, mRNA degradation (9), the oxidative stress response (10), and proteasome activity (11). Inhibition of Tor signaling is especially notable as this would induce a rapid reduction in synthesis for many of the most highly transcribed genes in yeast.

Recent progress has suggested that both thiolutin and holomycin act as redox-sensitive Zn^2+^ chelators, providing an explanation for their diverse cellular effects (1,11–13). It has been demonstrated *in vitro* that thiolutin and holomycin can be reduced by strong reductants such as DTT or TCEP (1,11) and that reduced thiolutin and holomycin can chelate Zn^2+^ (11,12). The Zn^2+^ chelating activity explains multiple thiolutin induced phenotypes, including alteration of glucose metabolism and proteasome inhibition through inhibition of Zn^2+^-specific metalloproteins (7,11). In addition, two observations suggest a possible mechanism for the thiolutin-induced expression of oxidative stress response genes (10). First, thiolutin and holomycin appear to require reduction for their activity, and reduction of thiolutin could in turn oxidize the cellular reductant or reducing proteins (1,12). Second, reduced holomycin can be spontaneously oxidized by molecular oxygen, suggesting that holomycin may be a redox cycling compound (14,15) and cause accumulation of reactive oxygen species (ROS). Whether thiolutin can act as a redox cycler has not yet been tested, and the cellular reductant(s) for thiolutin or holomycin remain complicated. Earlier reports indicated that glutathione does not reduce holomycin *in vitro* (1).

Recently, levels of glutathione in the range of those *in vivo* have been shown to reduce multiple dithiolopyrrolones *in vitro* and to activate Zn^2+^-chelating activity, consistent with these natural products being pro-chelators requiring reduction (16). Together, the Zn^2+^ chelating and redox activities are consistent with previously observed diversity in thiolutin-induced phenotypes, though transcription inhibition has not been easily explained. Both thiolutin and holomycin have been observed to be inert against RNAPs under conditions where either should be able to chelate Zn^2+^. Furthermore, it is not clear how Zn^2+^ chelation would be expected to inhibit transcription. In light of these negative results, transcription inhibition by dithiolopyrrolones was concluded to be a secondary outcome of inhibition of a distinct target, such as the proteasome. For example, thiolutin elicits removal of zinc from Rpn11, an essential deubiquitinase required for the proteasomal processing of ubiquitinated substrates (11,12).

There is extensive evidence for thiolutin inhibition of transcription *in vivo* in multiple species (17–19), though from *in vitro* studies it has been concluded that it inhibits prokaryotic and eukaryotic RNAPs differently. Thiolutin inhibited all three partially purified yeast RNAPs *in vitro* (20), but failed to inhibit all tested prokaryotic RNAPs *in vitro* (18,19,21). Similar RNAP inhibition *in vivo* with lack of inhibition *in vitro* has been observed for holomycin (12,22). The recently reported lack of thiolutin inhibition for all three fully purified yeast RNAPs has added further confusion (11). Therefore, our current state of understanding is that thiolutin and holomycin do not appear to inhibit prokaryotic or fully purified eukaryotic RNAPs, suggesting indirect modes of *in vivo* transcription inhibition. However, without comprehensive assessment of the experimental parameters for different transcription assays, it seems difficult to rule out direct mechanisms of action for thiolutin on eukaryotic RNAPs, as recent work does not satisfactorily explain why thiolutin inhibited all three partially purified yeast RNA polymerases *in vitro* (20), and how thiolutin inhibited RNAPs prior to template addition, but did not inhibit the template DNA bound RNAPs (20). This critical order-of-addition requirement for treatment prior to DNA addition is reminiscent of RNAP switch region inhibitor behavior (23,24) and is consistent with inhibition of a specific and early step in transcription, in contrast to Zn^2+^ chelation.

We can imagine a number of possibilities to reconcile the existing data. First, thiolutin may target a secondary protein, which is present *in vivo* and in partially purified RNA polymerase fractions (20) but not others (11,19,21). Second, as predicted (11,12), thiolutin may act as a prodrug that is activated *in vivo* and under the *in vitro* conditions of the early report (20) but not in assays used by others (11,19,21). To further investigate the mode of action of thiolutin, we have undertaken multiple approaches to screen for thiolutin resistant and sensitive mutants, as similar studies have contributed to understanding of modes of actions of many compounds (25–27), including holomycin (12). We identify a drug efflux pump functioning in thiolutin resistance and demonstrate that thiolutin treatment appears to induce oxidative stress. We propose that thiolutin likely induces the oxidative stress via oxidizing thioredoxins and possibly also through redox-cycling. In addition, we confirm that reduced thiolutin directly interacts with Zn^2+^ *in vitro*, consistent with the recent reports on holomycin and thiolutin using different approaches (11,12). We also suggest that thiolutin alters cellular Zn^2+^ homeostasis in yeast similar to holomycin’s effects in *E.coli* (12) but also find complex interactions with Co^2+^, Cu^2+^ and Mn^2+^ that are distinct from those of 1,10-phenanthroline, another class of Zn^2+^ chelator that inhibits transcription *in vivo* but has been described as an inhibitor *in vitro* through a Cu-chelate (28).

Finally, we have revisited *in vitro* biochemical studies with thiolutin and Pol II. We find that thiolutin indeed inhibits Pol II but critically requires the presence of small amounts of Mn^2+^, providing an explanation for disparate results across studies. First, DNA bound Pol II is resistant to thiolutin inhibition, recapitulating Tipper’s early study (20). Second, though thiolutin appears to inhibit a very early step in Pol II transcription, when initiation is bypassed by use of specific nucleic acid templates, thiolutin-treated Pol II exhibits pause-prone, slow elongation. Third, high DTT reverses the thiolutin inhibition, suggesting involvement of disulfide bond or redox sensitive covalent chemistry in inhibition. We find that thiolutin appears to inhibit initiation *in vivo* and reduces Pol II occupancy near promoters across many yeast genes. However, the most strongly affected genes are ribosomal protein (RP) genes with ribosome biogenesis (RiBi) genes being less but still strongly affected, consistent with rapid inhibition of Tor pathway signaling and downregulation of Tor pathway targets (29,30). Elongation of Pol II on genes also appears altered in that Pol II does not appear to completely run off longer genes as it would if elongation were unaltered, or if sole effects of thiolutin were indirectly through distinct signaling pathways. We propose that thiolutin inhibits Pol II transcription directly *in vitro* through a novel mechanism distinct from known Pol II transcription inhibitors but *in vivo* effects are complicated by high levels of pleiotropy. We caution that thiolutin use *in vivo* should be restricted to experiments aiming at understanding its complex modes of action and use as a tool for transcription inhibition *in vivo* would be fraught with undesirable complexity.

## Material and Methods

### Yeast strains and reagents

Yeast strains and primers used are listed in Table S2. *YAP1* C-terminal tagging with EGFP and gene deletion strains were constructed as described (31,32). Chemicals were commercially obtained from the following: Cayman Chemical (Thiolutin), Gold Biotechnology (DTT, TCEP, 5FOA), Sigma (MnCl_2_), JT Baker (ZnCl_2_), BDH Chemicals (MgCl_2_).

### Variomics screens

Two separate pools of diploid Variomics libraries were gifts from Dr. Xuewen Pan: one for non-essential and the other for essential genes (33). The pooled diploid Variomics libraries were grown, sporulated and selected for haploid Variomics libraries as described (33). Haploid Variomics libraries were subsequently used for the genetic screens. To convert the Variomics libraries into plasmid-free deletion libraries, pooled Variomics libraries (diploid for essential genes and haploid for non-essential genes) were grown in liquid SC media and plated on SC+5FOA plates to select against the mutants with the *URA3* plasmids containing the Variomics mutations. The diversity of the libraries were confirmed by deep sequencing of barcodes (“uptags” and “downtags”) flanking the deletion cassettes.

For manual screening of variomics libraries, about 1-3×10^7^ cells were plated on SC+10µg/mL thiolutin plates, and the potential thiolutin resistant candidates were restruck out on SC+10µg/mL thiolutin plates to validate the resistance. The plasmids from the validated resistant candidates were subsequently recovered for Sanger sequencing to identify putative resistance-conferring variants. We subsequently tested dominance for the repeatedly isolated candidates. The recovered plasmids from all reproducibly isolated candidates were transformed into the wild-type (CKY457) strain, and thiolutin sensitivity for transformants was phenotyped and compared to the empty vector control.

For high-throughput screens, Variomics libraries were screened on plates, while deletion libraries were screened in liquid, as previously described (25,26,33), and three independent screens (biological replicates) were performed. For Variomics screens, each library was screened on SC-URA+DMSO, SC-URA+8µg/mL thiolutin or SC-URA+10µg/mL plates. Each biological replicate was screened on 9 plates (6×10^7^ cells/plate) for 3 days, and cells were scraped to screen for an additional set of 9 plates for 3 days (6×10^7^ cells/plate). Deletion libraries were grown in liquid SC media to 3×10^7^ cells/mL and diluted in SC+3µg/mL thiolutin (for haploid non-essential gene deletion library) or SC+4µg/mL thiolutin (for diploid essential gene deletion library) to grow for 20 generations. During the selection, the cells were diluted to 1×10^6^ cells/mL every five generations to maintain cultures in log phase. For both screens, yeast cells were pooled, and genomic DNA was prepared using a YeaStar Genomic DNA kit (Zymo Research) for subsequent PCR amplification of barcode regions. Amplicons were sequenced by illumina Hiseq2500 in rapid mode. The sequencing reads are deposited in the SRA under the Bioproject PRJNA725683.

### Bar-seq data processing

Sequencing reads are mapped to the re-annotated barcode sequences (25) using Bowtie2 (34) (version 2.2.4) with the -N flag set to 0 and the --no-unal flag to suppress unaligned reads for further analyses. Bowtie2 outputs were written into SAM format and further extracted using Samtools (version 1.3.1) (35). Barcode sequences shorter than 15nts or were mapped to multiple reference barcodes were discarded. On average, 98% of sequencing reads were uniquely mapped.

As with Robinson *et al.* (36), we filtered out barcodes with lower than 100 total reads across all samples, since low count barcodes across all conditions were likely from sequencing errors and did not exist. In addition, mutants with less than 20 reads in both treated and the corresponding untreated controls are further filtered out, to exclude large changes from a small amount of reads with low confidence.

Following Robinson *et al.* (36), we added 1 pseudo-count to each barcode count to avoid division by 0, and performed TMM normalization using the edgeR package (version 3.12.1) (37). For differential abundance analyses, we used the edgeR to compute mutant specific dispersions, p-value and log transformed abundance changes using exactTest function (default setting). The transformed data were subsequently subjected to differential abundance analyses and clustering analyses. Differential abundance analyses were performed independently for the uptag and downtag sequences, and reproducibility between uptag and downtag data was assessed with Pearson correlation in R. Hierarchical clustering of the thiolutin phenotypic profile to an existing drug response profile for yeast mutants, consisting of 3356 compounds (26) was performed in Cluster 3.0 (38), using centered correlation and complete linkage.

Mutants with significantly altered abundance (p<0.01) were subjected to gene-ontology (GO) analyses as previously reported (26). In brief, all three GO categories, including biological processes (BP), Molecular Function (MF) and Cellular components (CC), were included in our analyses. Following the previous report (26), BP and MF GO terms that are too specific (present in less than 5 genes) or too generic (present in greater than 300 genes) were excluded, although smaller GO term groups were allowed for protein complexes (more than 2 genes) and larger groups (greater than 300 genes) were allowed for cellular components. We computed the GO term enrichment analyses for the significantly resistant or sensitive mutants (p<0.01) using hypergeometric test in R (version 3.2.2, physer function, default setting). Raw counts and p-value for each GO term were reported.

A series of quality controls were performed on the reproducibility among biological replicates, consistency between the two barcodes on each gene deletion (uptag and downtag) for the same strain, common and distinct properties between Variomics and Deletion libraries following a well-established statistical analysis pipeline. A subset of the statistically significant resistant or sensitive mutants were further validated by re-constructing the deletion strain for individual phenotyping.

### Visualization of Yap1 localization under fluorescence microscopy

Microscopy experiments were performed as previously described (39). In brief, CKY2038 was grown to mid-log (1-2×10^7^ cells/mL) in SC liquid media and added to the ConA treated perfusion chamber gasket (ThermoFisher, 4 chamber: 19 × 6mm) for treatment at indicated conditions (SC+1% DMSO, SC+10µg/mL Thiolutin, SC+10µg/mL Holomycin or SC+0.4mM H_2_O_2_) under the fluorescence microscope. The procedure to prepare the chamber and the microscope setting were the same as previously described.

### Glutathione quantitation assays

WT yeast (CKY457) cultures were grown (in YPD at 30°C) until mid-log phase (∼2×10^7^ cells/mL), washed and resuspended in the indicated conditions (YPD+1% DMSO, YPD+10µg/mL Thiolutin or YPD+10µg/mL Holomycin). The suspended cultures were grown at 30°C for one hour, and 5×10^7^/mL cells were subject to glutathione (GSH) quantitation. The active and total GSH levels were quantified using the GSH detection kit from Arbor Assays (K006-F5) following the manual. The glutathione disulfide (GSSG) level was calculated by (Total GSH - Active GSH)/2.

### UV-Vis assays

Thiolutin reduction and Zn^2+^ chelation reactions were performed in 250µL reactions with 50µM Thiolutin, 50µM ZnCl_2_ and 100mM potassium phosphate buffer (pH=6.5), as previously described (1). Mn^2+^ chelation reactions and relevant controls were performed under almost identical conditions except for the 100mM Tris buffer (pH=8) to keep consistent pH with the *in vitro* transcription assays. Each reaction had 250µL final volume and was quantified in Nanodrop 2000c spectrophotometer using UV-transparent cuvettes.

### Growth, viability and canavanine resistance assays

Phenotyping of mutants using plate assays were performed as previously described, with indicated addition of chemicals (40,41). Yeast liquid growth curve assays with Tecan Infinite F200 plate readers were performed as previously described (39). For viability assays and canavanine resistance assays, WT yeast strain CKY457 was grown in YPD until mid-log (1-2×10^7^ cells/mL), washed and treated with the indicated conditions (YPD, YPD+3µg/mL thiolutin, YPD+5µg/mL thiolutin and YPD+10µg/mL thiolutin) for an hour. The viability was quantified by staining the yeast cells in 0.1% Trypan blue staining (GE Healthcare Life Sciences, Catalog Number SV30084.01) for 3 mins, and the viability was quantified by counting the fraction of unstained yeast cells under the microscope. At least 300 yeast cells were counted in each repeat. Canavanine resistance frequency was quantified by plating the culture onto YPD and SC-Arg+60µg/mL canavanine plates and dividing the number of canavanine resistant colonies by the number of viable colonies on YPD plates. Three experimental replicates were performed.

### Pol II transcription activity assays

Pol II was purified using a strain expressing Rpb3 tagged with tandem-affinity tag (TAP), following a procedure as previously described (42). For initiation experiments DTT was removed from the Pol II buffer prior to the assay using Zeba spin desalting column (Thermo Scientific), though in later experiments this was found to be unnecessary.

Standard ssDNA transcription assays were performed in the *in vitro* transcription buffer (20mM Tris-HCl pH8, 40mM KCl, 5mM MgCl_2_). Each reaction had ∼0.5 pmol Pol II, 2µg denatured sheared salmon sperm DNA as the template and 200µM NTPs except 9µM cold NTP for labeling with that species (UTP or CTP), 1µM radioactive α-^32^P UTP or CTP (800 Ci/mmol, Perkin-Elmer) as substrates. The concentration of thiolutin, reductants (DTT or TCEP) and MnCl_2_ varies among the experiments (indicated in each figure). In general, 5 µl pre-treatment reactions including DMSO or thiolutin (at 2X final treatment concentration, from 7.5 µg/mL to 60 µg/mL in DMSO), +/− MnCl_2_ at specified concentration, and DTT (1-2X molar equivalent to thiolutin) were composed and incubated for 5 minutes at room temperature. To these reactions Pol II was added (with or without pre-binding to DNA) in an equal volume of 2X transcription buffer, yielding 10 µl reactions in 1X transcription buffer for thiolutin treatment. These reactions were incubated for 20 minutes at room temperature. Transcription was initiated by addition of 5 µl DNA/NTP (or just NTP mix in case of pretreatment) and incubated at room temperature for 30 minutes then stopped with stop buffer (10M Urea, 5mM EDTA in TBE). DTT reversal of thiolutin inhibition was accomplished by adding 2 µl 60 mM DTT to 8 µl Pol II treatment reaction at the end of the 20 minute treatment, prior to addition of NTPs/DNA. Transcripts were separated from unincorporated α-^32^P NTPs on 10% acrylamide/7M urea/TBE gels and visualized with a Bio-Rad Pharos Phosphorimager or GE Typhoon Imager. In vitro transcription elongation assays were performed as previously described (42), with modifications of time in the figure. 5µl reactions comprising 5 pmol Pol II, 40 µg/mL thiolutin in DMSO or DMSO alone, 875 µM MnCl_2_, equimolar DTT to thiolutin, 1X transcription buffer, and 3.75 µg BSA were incubated for 20 minutes at room temperature and incubated with 2.5 µl annealed transcription scaffold as described. After 5 minutes, non-template strand was added for an additional 5 minute incubation. Thiolutin-treated Pol II+scaffold was then diluted with additional 1X transcription buffer, MnCl_2_, thiolutin, while untreated Pol II+scaffold was diluted with thiolutin containing buffer (made fresh) or control buffer with DMSO. After 20 additional minutes at room temperature, reactions proceeded exactly as described in (42).

### ChIP-qPCR and ChIP-seq

(*RPB3::3XFLAG::kanMX4*) subunit of Pol II was used for chromatin immunoprecipitation (ChIP). For in vivo elongation assays, Pol II occupancy over a constitutive promoter-driven reporter (*TEF1p::YLR454W*) was determined after treating the cells with thiolutin. Yeast were grown in SC media to mid-log (1X10^7^ cells/mL) and treated with 10 µg/mL thiolutin for 2, 4, 8 or 10 mins (ChIP-qPCR) or for 1, 2, 4, or 8 mins (ChIP-seq), cross-linked with 1% formaldehyde solution (1% formaldehyde, 10mM NaCl, 100µM EDTA and 5mM HEPES-KOH, pH 7.5) for 20 minutes and quenched by 3M glycine (final concentration 360mM) for 5 minutes. ChIP-qPCR was performed as described previously (39). In brief, cross-linked cells were disrupted by bead beating and cell lysate was centrifuged at 1500 rpm for 1 min at 4°C to remove cell-debris, followed by centrifugation of chromatin pellets, subsequent washing of pellets (twice) with 1 ml FA buffer (50 mM HEPES-KOH pH 7.5, 300 mM NaCl, 1 mM EDTA, 0.1% Triton X-100, 0.01% sodium deoxycholate, 0.1% SDS and 1 mM PMSF) at 14000 rpm for 20 min at 4°C. Chromatin for ChIP-seq was washed one additional time in special FA buffer lacking SDS and sodium deoxycholate. Chromatin pellets were sonicated using a Diagenode Bioruptor (ChIP-qPCR, 45 cycles – 3 × 15 cycles; 30 s ON/45 s OFF) or a Diagenode Bioruptor Pico (ChIP-seq, 20 cycles; 30 s ON/30-50 s OFF) and used for IP with anti-FLAG antibody (FLAG M2 magnetic beads, Sigma-Aldrich). For ChIP-seq, 1100 µg of chromatin material as determined by BCA assay (Thermo Fisher) was used for each IP with addition of 110 µg *S. pombe* spike-in chromatin from CKP025 (*h-ura4-D18 rpb3+::6gly-3Flag-ura4+*). Concentration of SDS in IP for ChIP-seq after addition of *S. pombe* chromatin was 0.1% and concentration of sodium deoxycholate was 0.01%. SDS and sodium deoxycholate were omitted from all washes for ChIP-seq IPs as they reduce efficiency of FLAG M2 antibody (MilliporeSigma). 10% of the total IP material was used as input. Input or immunoprecipitated DNA was purified by standard phenol–chloroform extraction and ethanol precipitated with pellet paint (MilliporeSigma) or glycoblue (ThermoFisher). Immunoprecipitated DNA and 10% of corresponding input DNA (1:10 diluted) were used for qPCR with SsoAdvanced Universal SYBR Green supermix (Bio-Rad) using CFX 96 (Bio-Rad). Pol II fold enrichment was determined by the delta CT method.

### ChIP-seq library preparation

Libraries for ChIP-seq were prepared with NEBNext Ultra II DNA Library kit (NEB) according to the manufacturer’s instructions with the following modifications. Starting DNA for ChIP or input samples was 10 ng. Index primers were from NEBNext Multiplex Oligos for Illumina Dual Index Primer Set 1 and were diluted 1/2.5 before use. Library amplification during indexing amplification was between 6-8 cycles Samples were sequenced (paired-end) on an Illumina NextSeq 2000 P2 flow cell.

### ChIP-seq analysis

Paired-end raw fastq files were first trimmed for TruSeq3-PE-2 adaptors using Trimmomatic 0.38 (43) to remove adaptor readthrough. FastQC was used for quality control analysis. Adapter-trimmed fastq were then aligned to a hybrid genome combining *S. cerevisiae* and *S. pombe* genomes using bowtie2/2.4.5 (34), allowing 1 mismatch and disallowing insert size > 1000 (‘bowtie2 -q -N 1 -X 1000 –x’). The hybrid genome was generated by combining *S. cerevisiae* genome version R64-3-1 with *S. pombe* genome 2018-9-4, after appending a suffix of “pom_” to *S. pombe* chromosomes to allow reads mapping to *S. pombe* to be filtered later. Sam files generated from bowtie2 alignment were then converted to bam, which were then sorted and indexed using bamtools. Duplicates were then removed by picard 2.8.12, followed by indexing of newly generated bam files. Subsequent to duplicate removal, reads aligned to hybrid genome were segregated into separate species-specific (*S. cerevisiae* or *S. pombe*) input or IP bam files (‘samtools idxstats <non_duplicated-hybrid.bam> | cut -f 1 | grep -v “pom_” | xargs samtools view -b <non_duplicated-hybrid.bam> > <output_S.ce.bam>’; ‘samtools idxstats <non_duplicated-hybrid.bam> | cut -f 1 | grep “pom_” | xargs samtools view -b <non_duplicated-hybrid.bam> > <output_pombe.bam>’). Bams were then indexed using samtools/1.14 (35). This segregation of species-specific reads allows counting species specific reads in both input and IP bam files for normalization factor quantification. Total number of unique reads (samtools flag -F 260) per bam file of *S. cerevisiae* and *S. pombe* were counted for both input and IP data for calculating normalization factor combining input and spike-in reads, using an R script. The normalization factor was estimated following method of : (INPUT_Sp_Counts/100000000)/((Sp_Counts/100000000) * (INPUT_Sc_Counts/100000000)) (44). These factors were then used to scale the *S.cerevisiae* IP bams to generate scaled or normalized bigwigs using bamCoverage function of deeptools/3.3.0. Wiggletools/1.2.11 (45,46) was then used to generate mean bedgraphs of three replicates which were then converted to mean bigwigs using bedGraphtoBigWig of kentutils/v.370 (47). Differential bigwigs were generated by log2 operation function of bigwigCompare under the deeptools/3.3.0 package which were then used for generating heatmaps (46).

### 1H-NMR characterization of Thiolutin reduction and metal binding

Thiolutin stock solution for NMR experiments was freshly prepared by dissolving thiolutin in DMSO-d6. Thiolutin reduction study was performed using freshly prepared DTT in an NMR tube at the final sample concentrations of 1.5mM thiolutin, 7.5mM DTT, 15mM Tris (pH=8.0) and 10% D2O. 1H-NMR spectra were collected immediately using 500MHz Bruker Avance II spectrometer at 25 °C. For Zn^2+^ and Mn^2+^ chelation experiments, Zn^2+^ or Mn^2+^ were added at a final concentration of 7.5mM 1 minute after thiolutin reduction by DTT and were incubated at room temperature for 2 minutes before the acquisition of 1H-NMR spectra.

## Results

### Thiolutin or reduced thiolutin alone fail to inhibit fully purified Pol II in vitro

Recent publications suggest that thiolutin does not directly inhibit Pol II, in contrast to classic in vitro transcription experiments demonstrating thiolutin inhibition of partially purified yeast extract fractions containing Pol I, II, or III (20). We set out to understand this discrepancy by first examining thiolutin inhibition of purified Pol II. Given that thiolutin and holomycin activities could be sensitive to reduction of the disulfide bond by DTT (1,11,12) (Figure 1A), we removed DTT from the in vitro transcription reaction and performed thiolutin treatment with or without molar equivalents of DTT (Figure 1B). Under the buffer conditions employed here, we failed to observe thiolutin inhibition of Pol II, with DTT having no effect (Figure 1B). While these results are consistent with assertions that thiolutin does not directly target Pol II, they might similarly be explained by the absence of a cofactor or subtleties in experimental conditions between *in vitro* assays. We therefore set out to screen for potential co-factors while these biochemical assays are revisited below.

**Figure 1.**
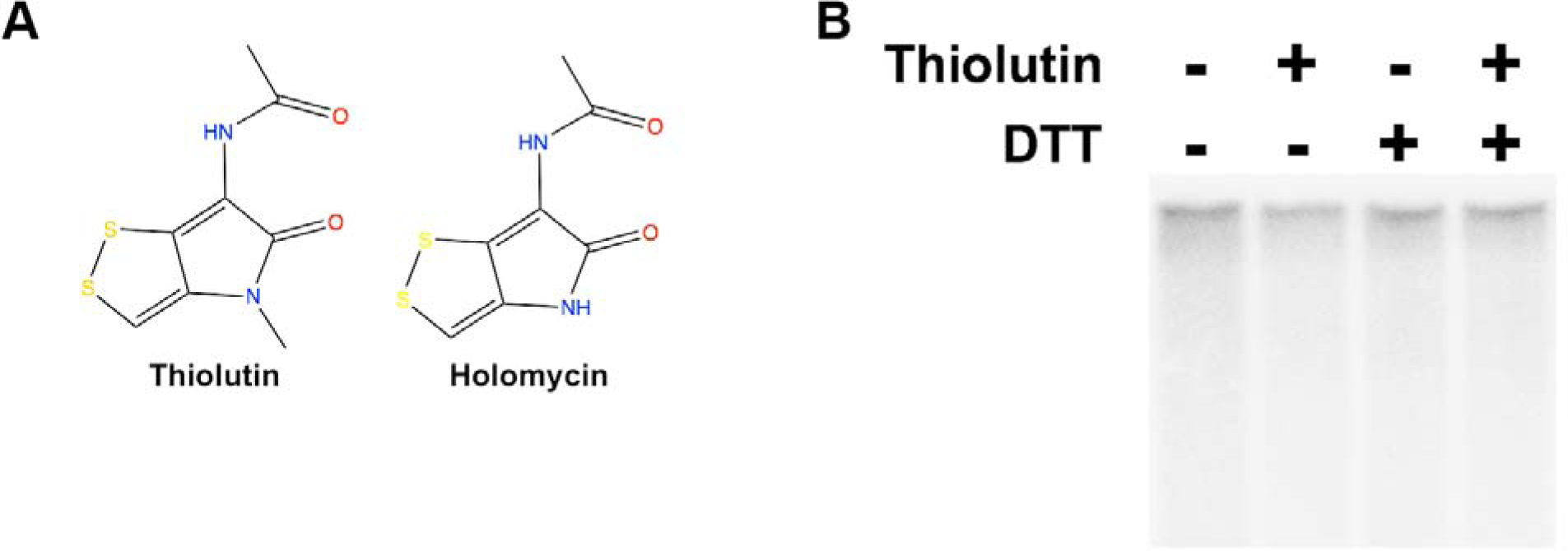
Thiolutin or reduced thiolutin alone fails to inhibit purified yeast Pol II *in vitro*. (A) Thiolutin and holomycin structures. (B) Thiolutin, or thiolutin treated with equimolar DTT, show no inhibition of Pol II transcription *in vitro*. Transcription activity assay was performed with ssDNA as the template and NTPs, including ^32^P labeled UTP. The transcribed ^32^P containing RNA was separated from free ^32^P UTPs on 10% polyacrylamide gels for visualization. Experiments were performed with at least three times, and a representative experiment is shown.

### Three independent genetic screens for thiolutin resistant and sensitive mutants

To gain insights into factors controlling the cellular response to thiolutin, we performed three independent genetic screens for modifiers of thiolutin sensitivity (Figure 2A, Figure S1A). First, we performed a conventional forward genetic screen for thiolutin resistant mutants through UV mutagenesis followed by thiolutin resistance selection. The causal mutations for thiolutin resistance were identified by bulk segregant analysis and whole genome sequencing (48) (Figure S1A). Second, we screened a yeast Variomics library for individual thiolutin resistant candidates (33). The Variomics library contained random mutations for almost all yeast genes cloned into a CEN based plasmid and expressed in strains with the relevant endogenous gene deleted. The CEN based plasmid used is known to propagate at 1-2 copies/cell and could result in increased gene dosage (49). Mutants from the forward genetics screen or those reproducibly isolated from the manual Variomics screens were re-tested for resistance (Figure S1B-C, Table S1, S2). Third, we performed a set of high-throughput screens using both pooled Variomics and deletion libraries using barcode-sequencing (Bar-seq) (25,26) (Figure 2A, Material and Methods), where the thiolutin resistant and sensitive mutants were identified by quantifying changes of mutant-linked DNA barcode frequency under thiolutin-treated or control conditions (26,36). Overall, our data exhibited excellent reproducibility (Figure S2) and revealed valuable common and distinct features between Variomics (putative gain-of-function (GOF) and loss-of-function (LOF) mutants combined) and deletion (LOF mutants) libraries (see Material and Methods). We have revealed contribution of multidrug resistance (Figs S1B-E, S3), oxidative stress response (Figs S1F-G, S5), multiple divalent metal homeostasis/trafficking pathways (Figs 2, 3, S7, S8) as well as the proteasome to thiolutin resistance (Fig S6), and we have further genetically dissected a subset of these pathways. These data collectively support that reduction and manganese or other metals appear to contribute to the thiolutin inhibitory activity, supporting a model that reduction and manganese are both required for thiolutin inhibition of Pol II in vitro, tested in subsequent biochemical experiments (Figs 4,5). Each of these pathways is discussed below.

**Figure 2.**
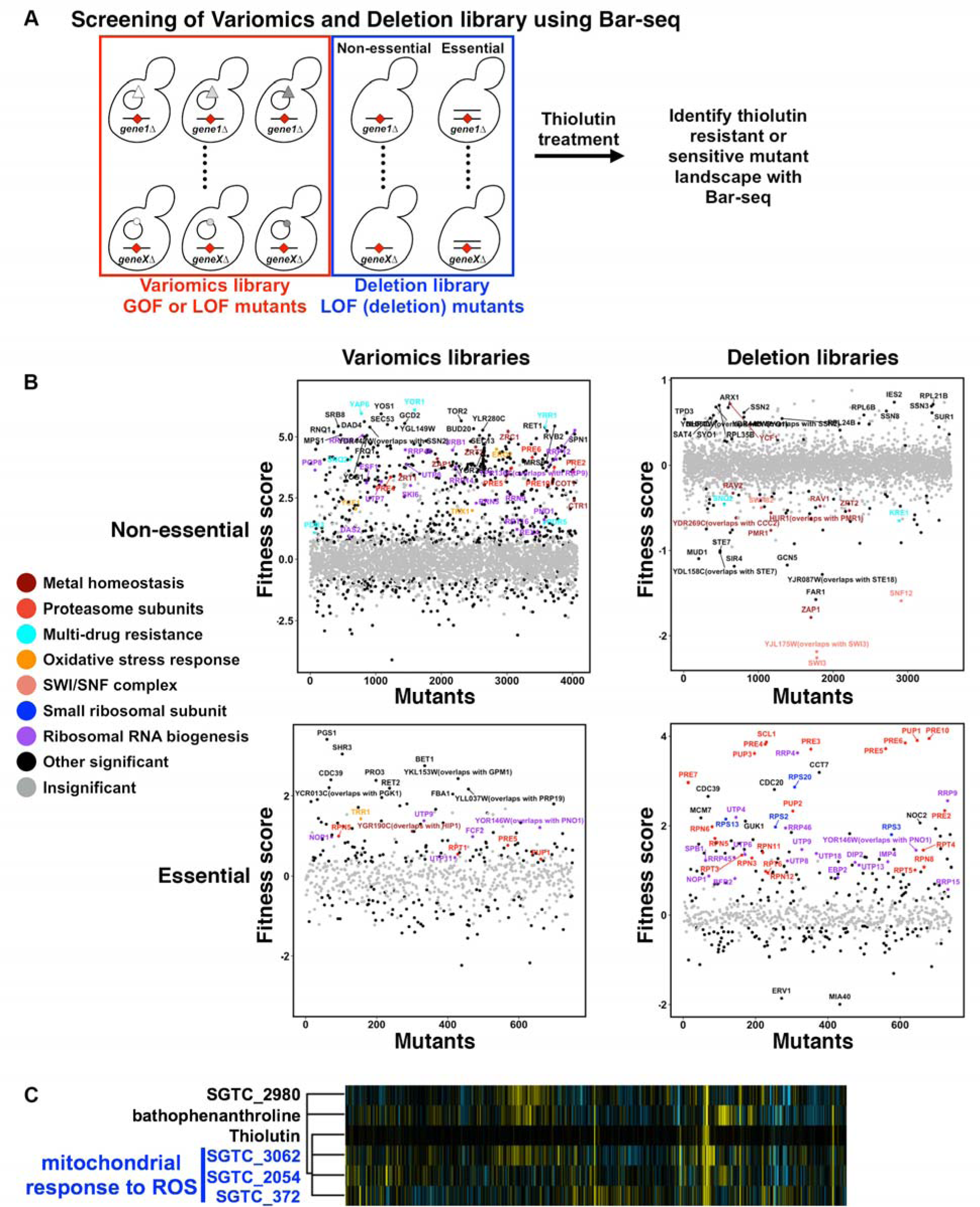
Mutations in diverse cellular pathways confer resistance/sensitivity to thiolutin. (A) Four libraries (Variomics libraries for non-essential or essential genes, Deletion libraries for non-essential or essential genes) were used for screening thiolutin resistant or sensitive mutants. Changes in abundance in the library was detected by deep sequencing of the PCR amplicon of the barcode region (Bar-seq). Variomics libraries consist of random gene variants and can be used to screen for both Gain-of-Function (GOF) and Loss-of-Function (LOF) alleles, whereas “Deletion” libraries consist of single-gene-deleted mutants that can be used to screen for complete LOF mutants. (B) Different thiolutin resistant or sensitive mutant classes are revealed in the Bar-seq based screenings of pooled Variomics and Deletion libraries. (C) Thiolutin induced phenotypic profile in the pooled deletion libraries co-clusters with drugs with signature of “mitochondrial response to ROS” (26) and bathophenanthroline, a metal chelator. 4683 deletion mutants’ responses (in columns) to 3356 compounds (in rows) were used for hierarchical clustering, but only the thiolutin closely correlated compounds are shown in this figure for clarity. Resistant strains: yellow; Sensitive strains: blue.

### Functional dissection of the multidrug resistance (MDR) and oxidative stress response (OSR) pathways in response to thiolutin

We identified thiolutin resistant strains harboring mutations in MDR and OSR genes from all three genetic screens. These results are perhaps not surprising given that yeast has well characterized MDR pathways that sense and pump out the toxic small molecules as a self-protection mechanism along with recent reports that thiolutin and holomycin activities require reduction in vitro (11,12). Reduction of thiolutin may itself oxidize cellular proteins and cause oxidative stress. In addition, it has been reported that reduced holomycin may act as a redox cycler, which could possibly generate ROS. Consistent with redox activity, thiolutin appeared to induce expression of several OSR genes (10). However, it has not been examined how thiolutin interacts with either MDR or OSR pathways. In light of our isolated thiolutin resistant mutants, we functionally dissected control of thiolutin sensitivity by these pathways. From both manual Variomics and forward genetic screens, we isolated multiple mutants in *YAP1* and *YRR1*, encoding transcription factors that have been implicated in OSR and/or MDR, respectively (Figure S1C). In addition, we also isolated mutants in the MDR efflux pump gene *SNQ2* from the manual Variomics screen. Finally, from high-throughput Bar-seq screens, we identified distinct thiolutin resistance and sensitivity in additional mutants linked to MDR transcription factors or efflux pumps (Figure 2B). Several lines of evidence suggested that the isolated thiolutin resistant MDR mutants were GOF mutations. First, isolated *yap1* mutations were clustered in C-terminal regions, where a variety of mutants are known to cause GOF (50–54). Second, isolated MDR mutants from the manual Variomics screen were dominant or functioned through increased dosage (overexpression) (Figure S3). Third, in Bar-seq screens, we found Variomics mutants in several MDR genes (*YAP1, YRR1, SNQ2*) conferred thiolutin resistance whereas deletion of the same genes either conferred sensitivity or had no strong effect (Figure 2B), consistent with these MDR Variomics mutants being different from the gene deletion mutants (complete LOF).

To determine the nature of identified MDR mutants from our screens, we constructed a series of MDR gene deletion mutants and tested their sensitivity to thiolutin (Figure S1D). We found that none of the tested deletion mutants was thiolutin resistant (Figure S1D). The lack of resistance for all the tested MDR deletion strains is consistent with the isolated thiolutin resistant MDR mutants from Variomics and forward genetics screens increasing MDR activity. In addition, *pdr1*Δ and *snq2*Δ conferred hypersensitivity to thiolutin (Figure S1D), consistent with the Pdr1 and Snq2 functions in promoting thiolutin resistance for WT strains. Together, we conclude that the MDR transcription factor Pdr1 is the major transcription factor promoting efflux of thiolutin from the cell, whereas neither Yrr1 nor Yap1 appeared to promote baseline resistance, given that *yap1*Δ and *yrr1*Δ did not confer thiolutin hypersensitivity. However, increased Yap1 or Yrr1 activities through mutations or dosage may increase efflux pump expression to promote thiolutin resistance, through their known linkage to many multidrug efflux pumps (55–57).

Holomycin did not inhibit yeast cell growth at low concentration (22), and we recapitulated these observations (Figure S4A). Given that structurally similar compounds may have different cell permeability and efflux efficiency, we asked whether the same set of MDR deletion strains conferred sensitivity to holomycin, but failed to observe any (Figure S4A). This observation suggested that the lack of holomycin sensitivity may not be due to selective efficiency of drug efflux, although we cannot rule out the possibility that thiolutin and holomycin may be transported by different efflux pumps.

In addition to MDR pathways, we also identified thiolutin-modulating mutations in OSR-related genes, including *YAP1* (discussed above) and genes in the thioredoxin pathway. Thioredoxins are a series of small anti-oxidant proteins that primarily function in reducing specific cysteines in client proteins (reviewed in (58) and references therein), and in turn are themselves reduced by thioredoxin reductases (Figure S1F). In yeast, there are two thioredoxin reductases (cytoplasmic Trr1 and mitochondrial Trr2) and three thioredoxins (cytoplasmic Trx1, Trx2; mitochondrial Trx3) (Figure S1F). Mutants in *TRR1* were reproducibly isolated from our manual Variomics screen (Figure S1B,E; Table S1). In addition, *TRR1\** and *TRX1\** plasmid bearing strains obtained from from high-throughput screening of Variomics libraries were resistant to thiolutin but not *trr1*Δ or *trx1*Δ were not (Figures 2B), suggesting that *TRR1\** and *TRX1\** plasmids in the Variomics library could be GOF due to specific mutation or increased dosage of the plasmid.

*TRR1* was found to be essential in large scale gene deletion analysis (59) but nonessential in classical genetic experiments (60–64). To understand the nature of *TRR1* function in thiolutin resistance/sensitivity, we constructed a *trr1*Δ strain to directly test its thiolutin sensitivity (Figure S1G). Interestingly, *trr1*Δ conferred hypersensitivity to thiolutin (Figure S1G), consistent with Trr1 activity antagonizing thiolutin and specific *TRR1\** mutants or increased Trr1 dosage enhancing this antagonism. The Trr1 function in antagonizing thiolutin also suggests that our isolated *yap1* alleles could confer thiolutin resistance through promoting *TRR1* expression, among other mechanisms.

We conceive of *trr1*Δ hypersensitivity to thiolutin in two ways. It could be due to loss of Trr1 function in directly counteracting thiolutin activity or the accumulation of oxidized thioredoxins, or both. The observed *TRX1\**-linked thiolutin resistance in high-throughput Variomics screens suggested critical functions of thioredoxin(s) in thiolutin resistance downstream of *TRR1*. Therefore, we examined thioredoxin deletions directly for thiolutin resistance. We found that the *trx1*Δ, *trx2*Δ, or *trx1*Δ *trx2*Δ mutants did not confer thiolutin sensitivity (Figure S1G). In contrast, *trx1*Δ *trx2*Δ conferred thiolutin resistance, and further, suppressed thiolutin hypersensitivity of *trr1*Δ (Figure S1G), consistent with the hypothesis that *trr1*Δ hypersensitivity to thiolutin is due to accumulation of oxidized thioredoxins. The observed *trx1*Δ *trx2*Δ thiolutin resistance reveals critical requirement of Trx1 and Trx2 for thiolutin activity *in vivo* and suggests that Trx1 and Trx2 may function either directly or indirectly in thiolutin reduction. However, further experiments are needed to test the direct thiolutin reduction by Trx1 or Trx2.

Thiolutin resistance conferred by *yap1* alleles and thioredoxin mutants suggests possible connections to OSR. Therefore, we set to test the roles of two additional OSR genes, *TSA1*, encoding a thioredoxin peroxidase, and *SOD1*, encoding a cytosolic copper-zinc superoxide dismutase, in thiolutin resistance. Interestingly, *sod1*Δ, but not *tsa1*Δ, conferred hypersensitivity to thiolutin (Figure S1G). Given that holomycin is a known redox cycler, we asked if simple addition of H_2_O_2_ could mimic the thiolutin effect in tested thioredoxin and OSR gene deletion mutants (Figure S1G). Thiolutin sensitivities in tested strains were distinct from H_2_O_2_ sensitivities, suggesting distinct effects on cellular function. Finally, we tested if yeast could be sensitized to holomycin by the same set of mutants (Figure S4). Interestingly, *trr1*Δ and *sod1*Δ were slightly sensitive to 10 µg/mL holomycin (Figure S4), suggesting that the induction of oxidative stress could be general property of dithiolopyrrolone compounds and that yeast are at least somewhat permeable to holomycin.

### Thiolutin induces apparent oxidative stress

Multiple lines of evidence presented above indicated that thiolutin may induce oxidative stress and increased Yap1 function may promote thiolutin resistance. Yap1 functions by sensing the cellular oxidants through its C-terminal cysteines, causing it to translocate into the nucleus and up-regulating a series of OSR genes to counteract the cellular oxidants upon oxidative stress (50–54). Therefore, nuclear localization of Yap1 is in general a sign of increased Yap1 function and OSR (50,53,54). To test whether thiolutin treatment caused nuclear translocation of Yap1, we fused *YAP1* to sequence encoding EGFP on the Yap1 C-terminus, and monitored Yap1::EGFP localization after thiolutin treatment. Thiolutin indeed induced Yap1::EGFP nuclear localization, consistent with induction of oxidative stress, although the thiolutin-induced Yap1::EGFP nuclear translocation appeared to be slower and weaker than that induced by H_2_O_2_ (Figure S5). In addition, thiolutin depleted total glutathione within an hour of treatment (Figure S6A). However, thiolutin depleted both active GSH and GSSG, a behavior similar to glutathione synthesis defective mutants but distinct from several well-known oxidants such as H_2_O_2_ and menadione (65–67). In contrast, holomycin effect weak at best on Yap1 localization and did not affect cellular glutathione levels (Figure S6A), consistent with the lack of growth inhibition by holomycin and distinct activities or trafficking in yeast between the two structurally similar compounds.

Previous studies suggested that the reduced holomycin was spontaneously re-oxidized when exposed to air (1), consistent with the behavior of redox-cycling compounds. To test if thiolutin exhibited similar behavior, we monitored thiolutin reduction and re-oxidation by its unique UV absorbance due to the conjugated ene-dithiol groups (Figure S6B, C), as with the recent reports (1,11,12). Thiolutin could be reduced by DTT or TCEP, but not equimolar glutathione (Figure S6B), consistent with the behavior of holomycin (1). Reduced thiolutin could be also re-oxidized (Figure S6C), suggesting that thiolutin can act as a redox-cycler, similar to holomycin (1). Together, thiolutin induced the Yap1 nuclear accumulation and hypersensitivity of *sod1*Δ and *trr1*Δ mutant in vivo and appeared to be a redox cycler in vitro, consistent with the induction of oxidative stress.

DNA damage can be a consequence of redox-cycling and can lead to transcription inhibition and cytotoxicity (68,69). We asked if thiolutin treatment could cause DNA damage or decrease viability. We did not observe an increase in mutation rate as measured by resistance to canavanine, consistent with little or no effect on DNA damage rates, nor did we observe a viability defect over the period of treatment (Figure S6D, E). The lack of viability defect is consistent with an early report showing that thiolutin inhibition was reversible after several hours’ treatment (70). Together, we suggest that thiolutin is a redox cycler in vitro and possibly also in vivo but does not appear to inhibit transcription through widespread DNA damage.

### Additional cellular pathways are involved in thiolutin resistance

The high-throughput screens of pooled yeast Variomics and deletion libraries, compared to conventional forward genetics and manual Variomics screens, revealed several additional pathways in thiolutin resistance (Figure 2B). We isolated mutants involved in metal homeostasis, SWI/SNF complex, proteasome, ribosomal RNA biogenesis, and small ribosomal subunits (Figure 2B). Many observations from the Variomics screens were already independently confirmed from two conventional screens, direct deletion analyses, or the existing literature (Figure S1, Table S2). For example, deletion of SWI/SNF complex subunits *SWI3* and *SNF12* have been shown to confer sensitivity to a variety of divalent metals including Mn^2+^, Zn^2+^, Co^2+^ and Cd^2+^ (71–75), consistent with thiolutin perturbation of metal homeostasis (discussed more below). We further constructed 9 haploid (for non-essential genes) or heterozygous (for essential genes) deletion mutants to validate the observation from the bar-seq screens of the deletion libraries. We validated the observations in 7 out of 9 mutants, including one resistant and one sensitive mutant from the non-essential deletion pool, and 5 resistant mutants from essential heterozygous diploid pool (Figure S7). Among them, we validated the observed thiolutin resistance conferred by three strains heterozygous for deletions in proteasome subunit genes (*PRE10/pre10*Δ, *PUP1/pup1*Δ, *RPN5/rpn5*Δ) (Figure S7).

The reconstructed strains validated the surprising but clear observation that heterozygous deletion mutants in proteasome subunit genes universally conferred thiolutin resistance (Figure S7). This is surprising because thiolutin was recently demonstrated to be a proteasome inhibitor through Zn^2+^ chelation, and heterozygous proteasome subunit deletion mutants presumably may lead to reduced proteasome activity and would be expected to confer sensitivity to proteasome inhibitors. Interestingly, two recent reports observed similar paradoxical resistance to proteasome inhibitors in yeast strains and human cell lines with decreased level of the regulatory 19S proteasome subunits (76,77). In yeast and human, the fully assembled 26S proteasome consists of a 20S catalytic core and a 19S regulatory complex. It was demonstrated that decrease or transient inhibition of 19S proteasome subunits may in turn induce the level and activity of partially assembled 20S proteasome instead of the fully assembled 26S proteasome (77), resulting in a net increase of proteasome function. It was also recently suggested that specific reduced expression of 19S proteasome subunits induced an altered cellular state and altered the global transcriptome in response to proteasome inhibitors (78). However, our observation is distinct, because we observed decreases in both 19S and 20S proteasome subunits conferred thiolutin resistance. Whether the proteasome has increased level and activity in these strains has yet to be tested.

### Thiolutin alters divalent metal homeostasis

The recent detection of direct Zn^2+^ chelation activity of reduced thiolutin and holomycin explains multiple thiolutin-induced phenotypes, such as inhibition of proteasome and glucose metabolism (7,11,12). From the high-throughput screens, we observed distinct thiolutin resistance or sensitivity linked to diverse Zn^2+^ trafficking genes, including those encoding the master transcription regulator Zap1 and various Zn^2+^ transporters Zrt1, Zrt2, Zrc1 and Cot1 (Figure 2B). Variomics strains linked to *ZAP1, ZRT1, ZRT2, ZRC1* and *COT1* conferred thiolutin resistance whereas deletion mutants were either not resistant (*zrt1*Δ, *zrc1*Δ, *cot1*Δ) or hypersensitive (*zap1*Δ and *zrt2*Δ) (Figure 2B), suggesting that these Variomics mutants could be gain-of-function or dosage modifiers of thiolutin resistance.

We first constructed a *zap1*Δ strain and tested its sensitivity to thiolutin (Figure 3A). *zap1*Δ confers hypersensitivity to thiolutin, consistent with Zap1 function in cellular resistance to thiolutin and thiolutin alteration of Zn^2+^ homeostasis *in vivo*. Notably, *zap1*Δ also conferred hypersensitivity to holomycin (Figure S8A). This observation is consistent with the reports that altering zinc homeostasis is a conserved mechanism among dithiolopyrrolones, and that holomycin could deplete cellular Zn^2+^ or access other essential targets in yeast if zinc homeostasis were otherwise perturbed, and suggesting that yeast resistance to holomycin may be partially explained by efficient transportation of Zn^2+^ that counteracts holomycin. *zap1*Δ hyper-sensitivity to thiolutin and holomycin was much stronger than that to H2O2, suggesting the *zap1*Δ hypersensitivity to thiolutin and holomycin were distinct from general oxidative stress exacerbation by zinc deficiency (79–82). and likely to Zn^2+^ homeostasis defects. Finally, Zn^2+^ supplementation at high concentration partially suppressed thiolutin inhibition (Figure S8C). The lack of full suppression suggests that there may be other thiolutin-mediated defects in addition to Zn^2+^ chelation, although we cannot rule out that inefficient Zn^2+^ trafficking limits effective Zn^2+^ supplementation in the cell.

**Figure 3.**
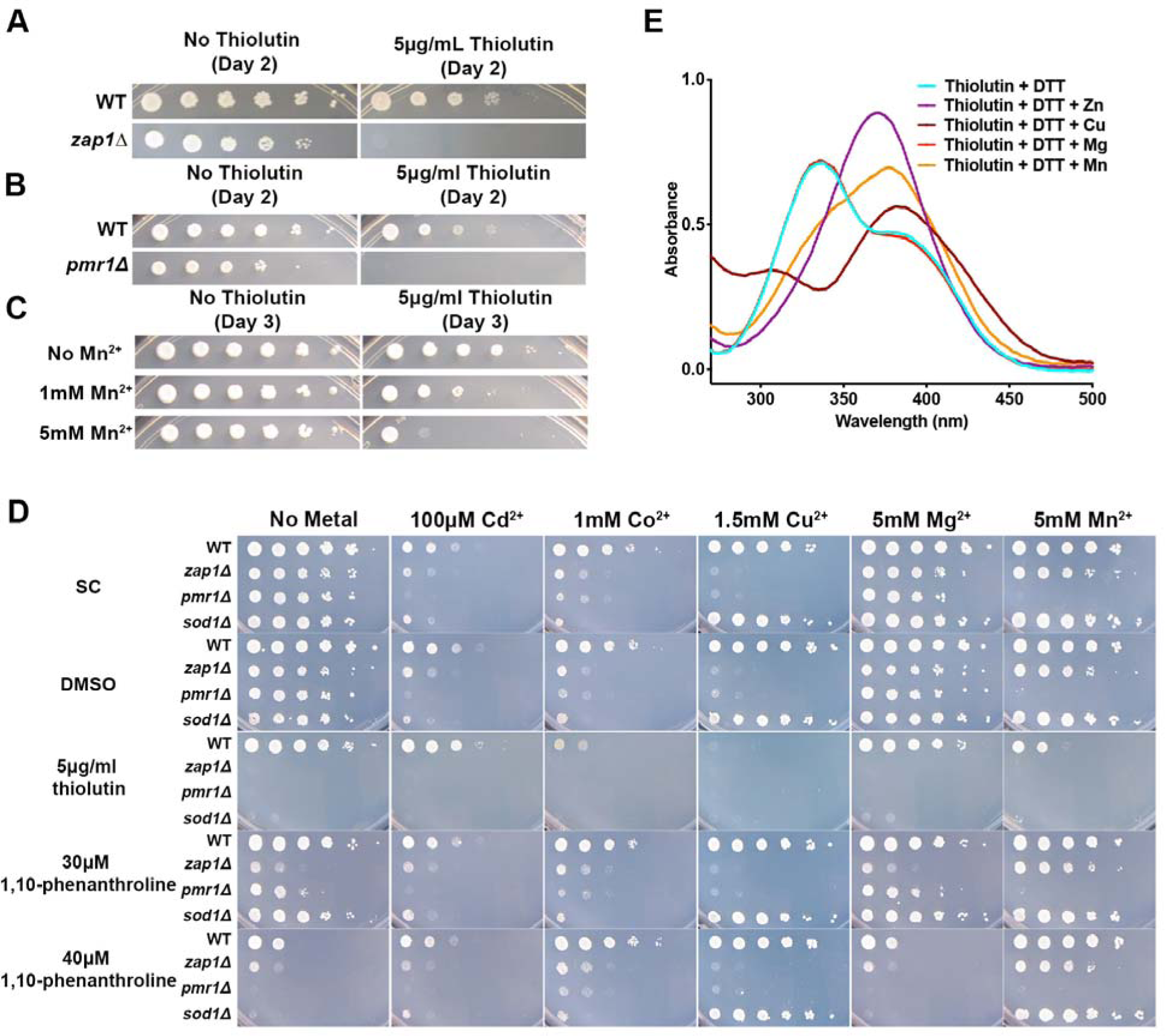
Divalent metals cations appear to alter thiolutin sensitivity *in vivo* and interact with reduced thiolutin *in vitro*. (A-D) 10-fold serial dilutions of yeast strains spotted onto different types of media to examine thiolutin sensitivity. (A) *zap1*Δ is hypersensitive to thiolutin. (B) *pmr1*Δ is hypersensitive to thiolutin. (C) Mn^2+^ supplementation exacerbates WT yeast sensitivity to thiolutin. (D) Distinct sensitivities of thiolutin and 1,10-phenanthroline to Co^2+^, Cu^2+^, and Mn^2+^ and differential interactions with gene deletions. (E) Reduced thiolutin was incubated with a selection of divalent metal cations and UV absorbance was analyzed. Zn^2+^, Mn^2+^, and Cu^2+^, but not Mg^2+^, interact with reduced thiolutin, as evident by alteration of the UV spectra of reduced thiolutin.

In addition to Zn^2+^ homeostasis mutants, we observed genes involved in Mn^2+^ trafficking (*PMR1, CCC2, VPS17, VPS25, SMF2*) linked to thiolutin resistance (Figure 2B). Interestingly, *PMR1* is a high affinity Ca^2+^ and Mn^2+^ ATPase that is required for Ca^2+^ and Mn^2+^ transport to Golgi and subsequent export (83,84). *pmr1*Δ has been shown to increase intracellular Mn^2+^ level and appeared to confer thiolutin sensitivity (Figure 2B), which we confirmed by reconstructing *pmr1*Δ mutant and plate phenotyping (Figure 3B, Figure S9A). In contrast, deletion of *SMF2*, a transporter for importing divalent metals including Mn^2+^, has been shown to cause intracellular Mn^2+^ depletion and conferred slight but consistent thiolutin resistance (Figure S8A). Together, the intracellular level of Mn^2+^ appeared to be positively correlated with thiolutin hypersensitivity, suggesting a mechanism distinct from Zn^2+^, where Zn^2+^ supplementation confers partial thiolutin resistance (Figure S8C). To rule out the possibility that this altered thiolutin sensitivity is due to indirect effect from interference of other divalent metals, we phenotyped wild-type yeast strain in the presence of thiolutin and Mn^2+^ supplementation. Indeed, we observed that Mn^2+^ dose dependently exacerbated thiolutin sensitivity in wild-type yeast, supporting the synergy between Mn^2+^ and thiolutin *in vivo* (Figure 3C).

In addition to *pmr1*Δ, we also confirmed the thiolutin hypersensitivity of *vps17*Δ and *vps25*Δ (Figure S9A). Vps17 and Vps25 are two vacuolar protein sorting genes and have been implicated in degradation of the Mn^2+^-importing protein Smf1, thus possibly causing intracellular accumulation of Mn^2+^ and conferring thiolutin hypersensitivity. However, *vps17*Δ and *vps25*Δ were not more sensitive to Mn^2+^ than WT in our strain background (Figure S9B) and we found that *smf1*Δ did not suppress the hypersensitivity conferred by *vps17*Δ or *vps25*Δ (Figure S9A), suggesting that other mechanisms may be involved in these two Vps mutants.

Given the contrasting Zn^2+^ and apparent Mn^2+^ modulation of thiolutin activity, and that metal homeostasis is complex with cross-talk between metals, we examined the intersection between divalent cations (Cd^2+^, Co^2+^, Cu^2+^, and Mg^2+^ to compare with Mn^2+^), thiolutin, and thiolutin-sensitive deletions (*zap1*Δ, *pmr1*Δ, and *sod1*Δ). We also compared thiolutin with 1,10-phenanthroline (1,10-pt), a chemically distinct Zn^2+^ chelator that can inhibit yeast transcription *in vivo* (85–87) (Figure 3D). Co^2+^, Cu^2+^, and Mn^2+^ sensitized yeast to thiolutin specifically but not to 1,10-pt. The tested deletion strains appeared more sensitive to thiolutin than to 1,10-pt at the concentrations tested. 1,10-pt sensitivity was highest for *zap1*Δ, consistent with 1,10-pt function as a Zn^2+^ chelator. We also observed distinct effects of different 1,10-pt levels and suppression of *sod1*Δ sensitivity to 40 µM 1,10-pt by Cu^2+^ and Mn^2+^. These results suggest that thiolutin and 1,10-pt are distinct, especially in relationships to metals other than Zn^2+^.

We tested whether reduced thiolutin chelates Mn^2+^ and Cu^2+^, while also confirming Zn^2+^ chelation. Consistent with two recent reports (11,12), Zn^2+^ caused a distinct UV shift of reduced thiolutin from ∼340nm to ∼370nm but did not change the UV spectra of non-reduced thiolutin (Figure 3E, Figure S9C). In addition, we found that Mn^2+^ and Cu^2+^ can also interact with reduced but not non-reduced thiolutin, in a manner likely similar to Zn^2+^ binding (Figure 3E). Consistently, 1H-NMR experiments confirmed that DTT caused chemical shifts consistent with reduction of the ene-disulfide in thiolutin, and further chemical shifts caused by diamagnetic Zn^2+^ and selective broadening caused by paramagnetic Mn^2+^ were consistent with interactions between the divalent metals and the thiolutin sulfhydryls (Figure S9, D and E). In contrast, Mg^2+^ did not bind to reduced thiolutin, distinct from the other tested metals (Figure 3E).

### Thiolutin, when activated by DTT and Mn^2+^, directly inhibits Pol II *in vitro*

Despite the progress in understanding thiolutin induced cellular phenotypes, whether and how thiolutin directly inhibits yeast Pol II *in vitro* remained unresolved. Multiple lines of evidence suggest possible involvement of divalent metals in this process. First, the co-clustering of thiolutin and bathophenanthroline induced phenotypic profiles suggests similar modes of action (Figure 2C). Bathophenanthroline is highly similar to 1,10-pt (Figure S10), which was suggested to require Cu^2+^ for direct transcription inhibition in vitro (28,88), though these compounds may have similar activities because they are also well-known Zn^2+^ chelators (89). Second, the direct interaction between reduced thiolutin and divalent metals (Zn^2+^, Mn^2+^ or Cu^2+^) suggests the possibility that thiolutin may function with multiple metal co-factors.

Many divalent metals inhibit RNAP activity (e.g. Pb^2+^, Zn^2+^, Cu^2+^, Be^2+^, Cd^2+^, Ca^2+^) (90), and additionally are tightly controlled in cells due to toxicity (*e.g.* Fe^2+^, Co^2+^), so may not participate in possible Pol II inhibition by thiolutin. Mn^2+^ does not inhibit RNAP but instead increases the activity of multiple RNA polymerases (including Pol II) (90). In addition, Mn^2+^ is readily available in yeast cells (0.04-2mM depending on the type of measurement while genetically shown to be at levels that likely can alter Pol II activity *in vivo*) (91–95) at a relevant range to the reported thiolutin inhibitory concentration (∼20µM in (20)). We note that in the original Tipper observations (20), their transcription buffer contained 1.6 mM Mn^2+^ as many classical transcription buffers included both Mn^2+^ and Mg^2+^ to maximize observed activity. In contrast, most other studies did not report Mn^2+^ in buffers (12,18,21,22) (One report appears to have a high concentration Mn^2+^ (10 mM) and possibly also reductant (19)). Therefore, we set out to test whether Mn^2+^ might participate in thiolutin-mediated inhibition of Pol II. Remarkably, Mn^2+^ addition to reduced thiolutin potently inhibited purified Pol II activity (Figure 4A), in sharp contrast to the activation of Pol II activity caused by Mn^2+^ by itself in the absence of reduced thiolutin. Our observation is consistent with the known stimulation of RNAPs by Mn^2+^ (90,96), but also suggests a potent Pol II inhibition by Mn^2+^ activated reduced thiolutin (hereafter termed as Thio/Mn^2+^) (Figure 4A, Quantification of thiolutin treatments and controls in Figure 4C,D).

**Figure 4.**
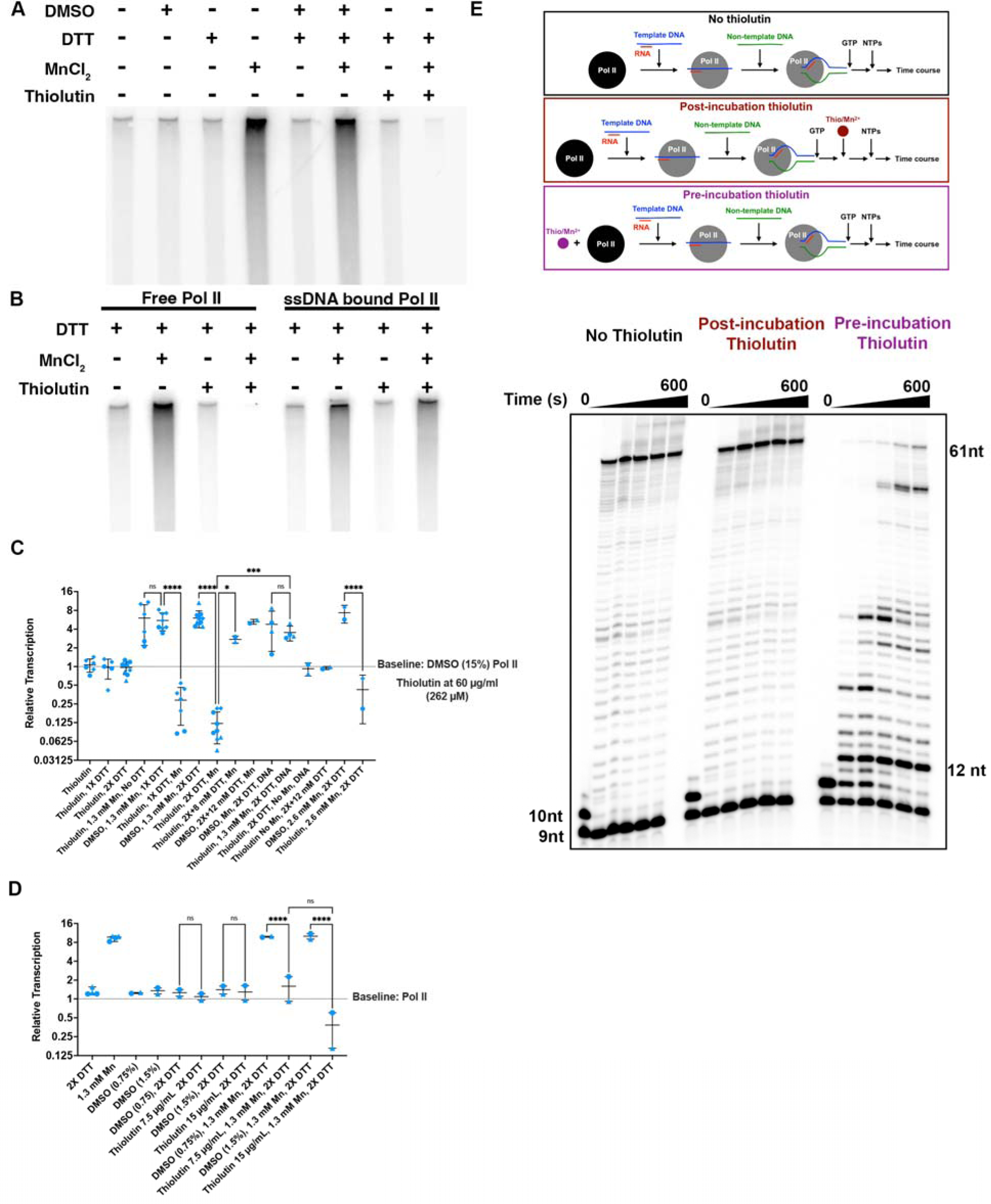
Apparent thiolutin/Mn^2+^ complex inhibits Pol II transcription *in vitro* and *in vivo*. (A) Thiolutin requires both DTT (reductant) and Mn^2+^ to inhibit Pol II transcription *in vitro*. A representative of multiple experiments is shown. Quantifications of this and a wide range of experimental conditions for thiolutin inhibition are shown in Figure S10. (B) Pre-binding to DNA renders Pol II resistant to Thiolutin/Mn^2+^ complex. This gel is representative of multiple independent experiments (Quantifications shown in Figure S10). (C) Quantification of thiolutin inhibition under a number of experimental conditions. Thiolutin (60 µg/mL) inhibition of Pol II requires DTT and Mn2+ and is blocked by pre-incubation of Pol II with DNA or with excess (12 mM) DTT added after Pol II treatment with thiolutin. All reactions are compared to baseline for individual Pol II in particular experiment, which is activity of Pol II plus vehicle (15% DMSO final). Different symbols indicate three independent Pol II purifications used in experiments. Note that variability in fraction of Pol II inhibited by thiolutin appears to track with Pol II purification (▴● Pol II consistently more inhibited than Ill). Error bars are mean +/− Standard Deviation. Statistical analysis for selected pairs is one-way ANOVA with multiple comparisons controlling for False Discovery Rate (0.01) (method of Benjamini, Krieger, Yekutieli (110)). **** = p< 0.0001, *** = p< 0.0002. * p< 0.0332. (D) Quantification of thiolutin inhibition at lower thiolutin/DMSO concentrations and additional controls. All reactions are compared to baseline for individual Pol II in particular experiment, which is activity of Pol II in absence of drug or vehicle. Different symbols indicate two separate Pol II purifications (symbols here match Pol II purifications in (C). Statistical analysis for selected pairs is one-way ANOVA with multiple comparisons controlling for False Discovery Rate (0.01) (method of Benjamini, Krieger, Yekutieli (110)). **** = p< 0.0001. (E) In vitro elongation assay for Thiolutin-treated Pol II. (Top) Schematic indicates three assembly pathways for making elongation complexes on nucleic acid scaffolds in the absence of thiolutin, incubation with thiolutin after elongation complex assembly, or before elongation complex assembly. (Bottom) Thio/Mn^2+^ inhibited Pol II can be directly assembled into a distinct and slow elongating complex, but only if Pol II is treated with thiolutin prior to complex assembly. Two experimental replicates were performed, and a representative replicate is shown. Duration times for incubation with NTPs were 0, 10, 30, 120, 300, and 600 seconds.

We next designed a series of experiments to investigate properties of the inhibited Pol II and nature of the inhibitory species. We first tested if order-of-addition among Pol II, DNA and Thio/Mn^2+^ was critical for the inhibition, since order-of-addition is often informative on the mode of transcription inhibition (Figure 4B, E). This is especially important given the observation of Tipper that order-of-addition was critical for thiolutin inhibition in that study (20). Consistent with the early observation, we found that the template binding protects Pol II from the Thio/Mn^2+^ inhibition (Figure 4B). Template protection of Pol II suggests Thio/Mn^2+^ inhibits Pol II DNA interaction or similarly early step in transcription, consistent with the behavior of transcription initiation inhibitor. Therefore, we tested if Thio/Mn^2+^ affected Pol II elongation on a transcription bubble template using an RNA primer, where transcription initiation is bypassed (Figure 4E). Surprisingly, Thio/Mn^2+^ altered, but did not block Pol II transcription elongation, inducing a highly pause prone Pol II elongation mode (Figure 4E). This was unexpected because most specific initiation inhibitors do not cause elongation defects, and DNA binding does not protect RNAPs from elongation inhibitors by their very nature. In addition, pre-assembled transcription elongation complexes were also resistant to Thio/Mn^2+^ (Figure 4E, bottom middle panel), validating the critical order-of-addition in the non-specific transcription assay (Figure 4B). Thio/Mn^2+^ inhibited Pol II was pause-prone and appeared to irreversibly arrest at specific template positions (Figure 4E, bottom right panel).

Whether thiolutin inhibits transcription initiation or elongation *in vivo* is unclear (17,21). In light of our *in vitro* results, we tested thiolutin-mediated Pol II transcription inhibition *in vivo*, first by monitoring Pol II occupancy on a long gene, *YLR454W* driven by the *TEF1* promoter, and then by Pol II ChIP-seq genome wide (Figure 5). We observed a specific decrease of Pol II occupancy by qPCR on the 5′ end of *YLR454W* within the first 2 minutes of thiolutin treatment, consistent with fast transcription initiation inhibition after thiolutin treatment (Figure 5). After 2 minutes, we also observed a relatively slower loss of Pol II from the template compared to previous experiments where transcription initiation was inhibited by other means, specifically by glucose shutoff or by carbon starvation of transcription from a *GAL1* promoter driven *YLR454W* reporter (39) (comparison with prior studies shown in Figure S11). Additionally, this decrease in Pol II occupancy at later time points does not fully cover the 3′ end as it does at *GAL1p::YLR454W* under the aforementioned inhibition conditions where the gene is clear of Pol II by 4-6 minutes (39). These results suggest that thiolutin may have additional non-immediate inhibitory effects on Pol II elongation *in vivo* and that thiolutin treatment is not functioning solely through altering cellular signaling (*e.g.* through inhibition of the Tor pathway (8)) (Figure 5A). To assess kinetics of effects on Pol II occupancy in response to thiolutin treatment genome wide, we performed ChIP-seq for a FLAG-tagged Rpb3 Pol II component in triplicate upon thiolutin treatment (1, 2, 4, 8 minutes), DMSO vehicle treatment (8 minutes), or control cells (Figure 5B-D, Figure S12). Our ChIP-seq protocol includes spike-in of Rpb3-FLAG chromatin from *S. pombe* allowing for determination of global effects (Figure 5B). Our experiments showed good reproducibility across replicates (Figure S12A,B)) with a clear trend of increased effects over time of thiolutin inhibition. We find evidence of broad decrease in Pol II occupancy genome wide (Figure 5C). We observe a continuum of effects for highly expressed genes where ribosomal protein (RP) genes are affected more strongly, then ribosome biogenesis genes (RiBi), then other genes (Figure 5D, 5E). These results are consistent with reported inhibition of the Tor pathway upon thiolutin treatment but there also more widespread, but muted decreases in Pol II occupancy. Consistent with our observation at *TEF1p::YLR454W*, metaplots for all genes in the top 40% of Pol II occupancy ≥1.5kb in length show an initial decrease in Pol II occupancy at the 5′ end upon thiolutin treatment (Figure S12C). The most striking effects of thiolutin on Pol II occupancy are those that occur with rapid kinetics, e.g. decreased occupancy on highly expressed genes, specifically of the RP and RiBi classes, and rapid increased occupancy of a small subset of genes. These classes of effect like involve in Tor signaling inhibition (8,29,30,97) and stress response activation (11,85), respectively. The more muted but widespread effects and potential slowing of elongation could be consistent with additional, direct or indirect effects of thiolutin that occur with reduced kinetics (see Discussion).

**Figure 5.**
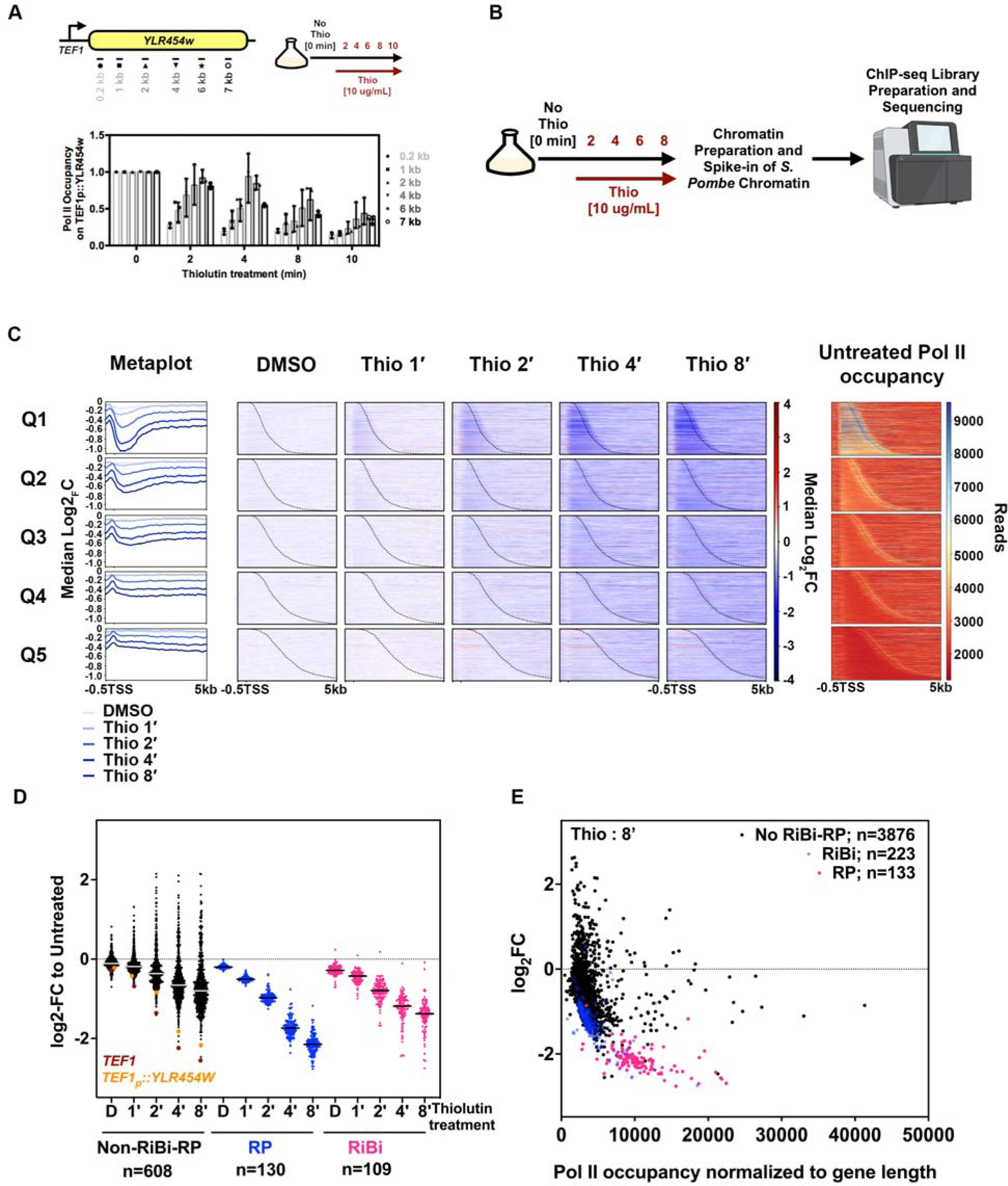
Thiolutin reduces Pol II occupancy genome wide. (A) Thiolutin inhibits Pol II occupancy at a *TEF1* promoter-driven *YLR454W* reporter *in vivo.* Schematic indicates positions of PCR primers for chromatin immunoprecipitation-quantitative PCR (ChIP-qPCR) for a FLAG-tagged Rpb3 subunit of Pol II at a reporter gene representing a long transcription unit. Thiolutin (10µg/mL) was added to cells for indicated times prior to crosslinking for ChIP. Three experimental replicates were performed, and error bars represent standard deviation of the mean. (B) Schematic of Pol II ChIP-seq experiment thiolutin treatment for the indicated times (control sample with addition of vehicle DMSO was for 8 minutes treatment). (C) Genome-wide decreases of Pol II occupancy at genes separated into quintile based on normalized Pol II occupancy in untreated cells from highest (Q1) to lowest (Q5). Quintiles are ∼846 genes each. Genes are rank ordered from shortest to longest with annotated gene 3′-ends indicated with the dashed line. (D) The majority of genes show decreases in Pol II occupancy as determined by fold change in normalized Pol II occupancy per base within each gene. Genes separated by class indicate that ribosomal protein (RP) genes show greater Pol II decreases than do ribosomal biogenesis (RiBi) than do other highly expressed genes in Q1. (E) Decrease in Pol II occupancy (y axis) shows some correlation with initial occupancy levels (x-axis) with RP genes showing special sensitivity to thiolutin treatment.

To investigate the nature of the inhibitory species, we further examined thiolutin interaction with Mn^2+^. Interestingly, we observed continuous and reproducible changes of the UV spectra after Mn^2+^ was added to reduced thiolutin (Figure 6A), suggesting changing chemical species in the reaction. The peak of UV absorbance was shifted from 340nm (reduced thiolutin) to around 380nm in the first two minutes, with subsequent decrease in the 380nm peak as a third species at 300nm started to accumulate (Figure 6A). The reaction reached a relative stable equilibrium after 20 minutes. Our data revealed the dynamic change and two major species after Mn^2+^ addition to the reduced thiolutin but could not distinguish how either of these related to the inhibitory species. To test this question, we prepared Thio/Mn^2+^ and either freshly treated Pol II or incubated at room temperature for 20 minutes before treating Pol II (Figure 6B). Remarkably, we observed strong Pol II inhibition by freshly prepared Thio/Mn^2+^ but complete loss of Thio/Mn^2+^ inhibitory activity after 20 minutes (Figure 6B). This result is consistent with the 380nm species possibly contributing to Pol II inhibition yet being unstable in solution. In contrast to the window for inhibitory activity of Thio/Mn^2+^, immediately treated Pol II was inhibited for up to 50 minutes (20-minute incubation and 30-minute reaction time), suggesting that the Pol II inhibition was stable (Figure 6B).

**Figure 6.**
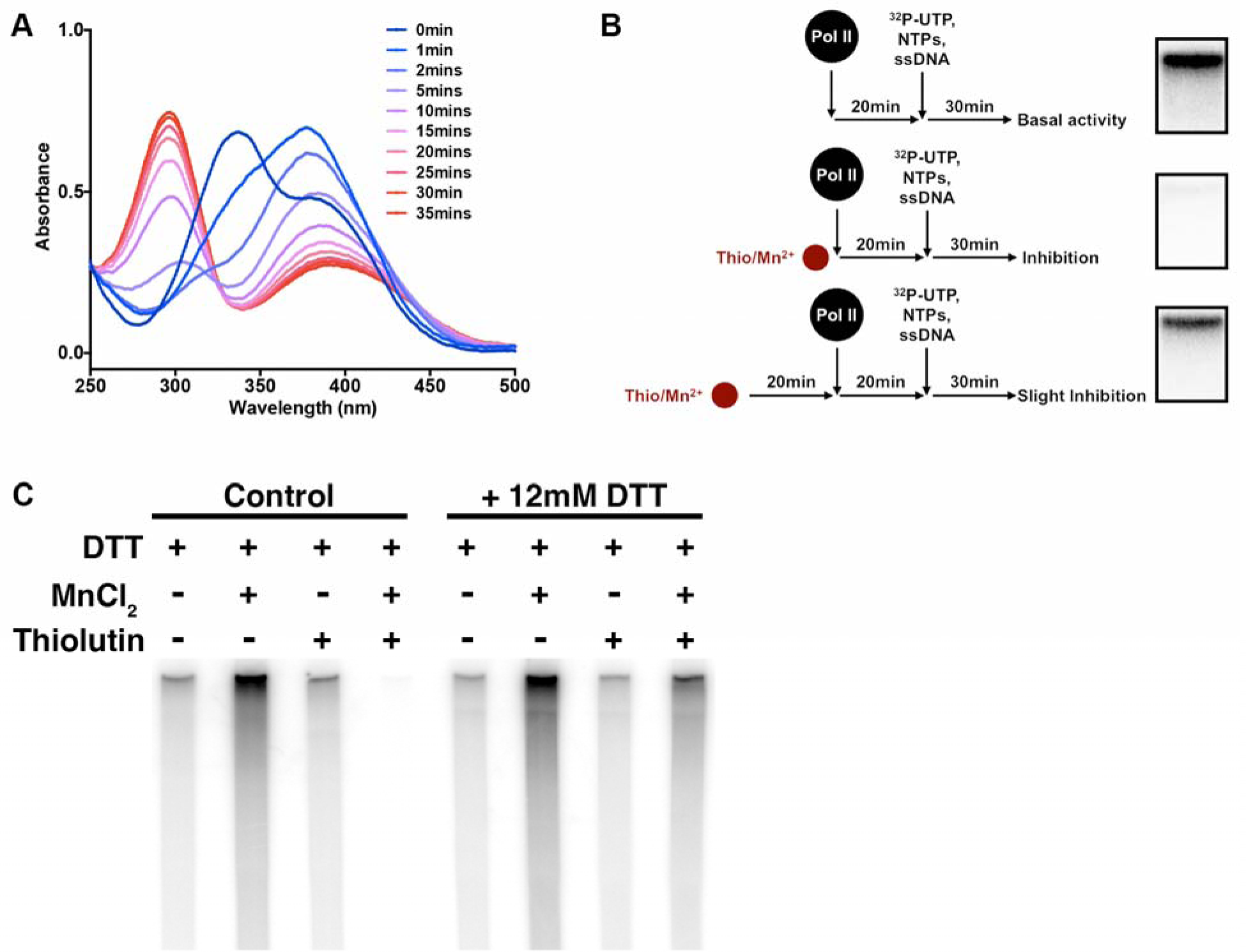
The apparent Thiolutin/Mn^2+^ complex is unstable in solution but Pol II is stably altered. (A) Time course of changing UV spectra of reduced thiolutin treated with Mn^2+^. Reduced thiolutin (blue, 0 min) was treated with equivalent molar Mn^2+^, and the UV spectra was acquired at different time points. Spectra over time was colored in a series of gradient from blue to red. Three experimental replicates were performed, and a representative replicate is shown. (B) Thio/Mn^2+^ lost inhibitory activity after 20 mins in solution though immediately treated Pol II remains inhibited. Two experimental replicates were performed and consistent. One replicate is shown. (C) Thio/Mn^2+^ can be reversed by excess DTT. Excess of DTT was added after 20 minutes of Pol II inhibition by Thio/Mn^2+^. DTT was at 12 mM during thiolutin treatment. Two replicates under these exact conditions were performed, and a representative replicate is shown. Results are representative of additional experiments performed under slightly different conditions.

We propose two models to reconcile the observation that Pol II stabilized the unstable 380nm inhibitory species. First, Pol II might stabilize the inhibitory species in a tight pocket that prevented further reaction into other inactive species. Second, Mn^2+^ might coordinate with reduced thiolutin to form a disulfide bond within Pol II or possibly a thiolutin adduct, as Cu^2+^ does to facilitate disulfide formation through cysteine sulfenylation in other proteins (98–100). If sulfenylation and disulfide bond formation were involved, the Thio/Mn^2+^ inhibition is expected to be suppressible by a high concentration of DTT. Indeed, we found that excess DTT abolished the inhibition, consistent with the involvement of reductant sensitive inhibition (Figure 6C).

## Discussion

Thiolutin is a well-known transcription inhibitor that has been used in mRNA stability studies, but its mode of inhibition has been complicated and unresolved. It has been demonstrated that reduced thiolutin and the related natural product holomycin chelate Zn^2+^ in vitro, and thiolutin chelation of Zn^2+^ could specifically inhibit diverse classes of metalloproteins (11,12). However, these previous reports failed to observe direct thiolutin inhibition of RNAPs, under conditions where Zn^2+^ chelation was permissible. Here we present three independent genetic screens for thiolutin resistant or sensitive mutants, providing a genetic basis for understanding thiolutin-altered cellular responses and reveal a direct mode of action against Pol II *in vivo*. We show that alterations in diverse cellular pathways can modulate cellular sensitivity to thiolutin. In light of our genetic results, we discovered that both reductant DTT and Mn^2+^ together activate thiolutin for direct inhibition of Pol II function, countering the recent narrative while confirming classic observations (11,20).

### Thiolutin inhibits Pol II through a novel mode of action

We propose that Thio/Mn^2+^ inhibits Pol II through a novel mode of action. The critical order of treating Pol II with thiolutin prior to template DNA binding is a stereotypical behavior for RNAP clamp inhibitors (23,24,101–104), which are distinct from inhibitors targeting the active site, NTP uptake, or RNA exit channels. In addition, we show that thiolutin inhibits Pol II elongation on an initiation-bypassing transcription bubble template, distinct from all the three characterized *E. coli* RNAP inhibitors that lock the clamp in the closed state (Myxopronin (Myx), Corallopyronin (Cor), and Ripostatin (Rip)) that do not inhibit RNAP elongation (23,105). It should be noted that Lipiarmycin (Lpm), an inhibitor that appears to lock the clamp in the partially or fully open state (24), has never been tested on a bubble template. Therefore, thiolutin behaves differently from Myx, Cor and Rip, but further experiments are in need to test whether thiolutin behaves similarly to Lpm. Finally, the exact thiolutin binding site on Pol II remains unclear, and our data cannot rule out the possibility that thiolutin may inhibit Pol II regions other than clamp-controlling switch regions.

### Diverse cellular pathways modulate thiolutin multiple modes of action

Our genetic screens reveal the contribution of diverse cellular pathways in thiolutin sensitivity. Many of these pathways can be reconciled in a unified model based on current understanding of thiolutin mode of action. In the cell, the thiolutin intra-molecular disulfide bond appears to be reduced, and we propose thioredoxins Trx1 and Trx2 directly or indirectly contribute to thiolutin reduction (along with glutathione) or are responsive to thiolutin oxidation of other proteins, thereby promoting thioredoxin oxidation. Reduced thiolutin subsequently chelates Zn^2+^ to inhibit multiple metalloproteins (*e.g.* Rpn11 and conferring block to proteasomal degradation of substrates), affects Zn^2+^ homeostasis, interacts with Mn^2+^ to inhibit Pol II transcription (and possibly Pol I and III), or become re-oxidized by molecular oxygen. Mn^2+^, Co^2+^, and Cu^2+^ also sensitize cells to thiolutin treatment while 1,10-pt does not, suggesting these interactions may not be caused by Zn^2+^ starvation. The reduction or redox cycling of thiolutin may induce the observed Yap1 nuclear localization and oxidative stresses. In addition, we have shown that reduced thiolutin may also chelate other divalent metals such as Cu^2+^ (but not Mg^2+^), but whether other thiolutin-metal complexes have additional activities remain to be further tested.

### Thiolutin should not be used as a tool for transcription inhibition *in vivo*

Our work underpins the caveats for using thiolutin as a transcription inhibitor to investigate other cellular processes, as suggested by previous studies (9,11,12). As a routinely used transcription inhibitor to study mRNA stability, it was shown that thiolutin itself inhibits mRNA degradation and complicates the quantitation of mRNA half-life at higher doses (9). The rapid kinetics of Pol II occupancy changes we observe for RP genes are consistent with a potential immediate perturbation to Tor signaling. Note that a previous study only examined Tor pathway response at 10 minutes of thiolutin or 1,10-pt treatment, whereas our work suggests potential effects would be nearly immediate. RiBi genes are also downstream of the Tor pathway (30,97) are strongly affected for Pol II occupancy. However, most other genes of similar expression level to RiBi show similar decreases in Pol II occupancy (Figure 5E). Net decreases in occupancy are widespread (Figure 5D, E) but a small subset of genes is activated by thiolutin treatment with rapid kinetics (Figure S12B). Activation of this subset is consistent with an immediate stress response that precedes any potentially slower, direct effects of thiolutin (11,85,106). Our observation that elongation of a long reporter gene, *TEF1p::YLR454W*, appears slow upon thiolutin inhibition of initiation is striking and points to effects that require careful dissection in future experiments. Our work here and prior work discussed above suggest that thiolutin is multifunctional and has pleiotropic effects.

### Thiolutin mode of action may reveal insights into Pol II pausing

The thiolutin induced pause (and arrest) pattern *in vitro* appears to be highly specific and position-dependent, and it will be interesting to further investigate the pause sequence preference and its possible connection to a specific inhibited Pol II conformation. As discussed above, biochemical properties of thiolutin inhibited Pol II most closely resembles clamp inhibition by a locked non-closed state, possibly a partially or fully open state. It is tempting to hypothesize that the thiolutin-induced pause prone Pol II is linked to a specific Pol II conformation. This hypothesis is also consistent with the observations in *E. coli* RNAP that paused elongation complexes appear to correlate with open clamp and TL states, which can be reversed and possibly regulated by elongation factor RfaH (107,108). It has been proposed that this RfaH (a homolog of Spt5 in eukaryotes) pause suppression through the RNAP clamp may be a conserved regulatory mechanism for RNAP elongation in all domains of life (109). Further characterization of thiolutin inhibited Pol II may reveal additional insights into this process in eukaryotes.

## Supporting information

Table S1 Variomics Candidates

Table S2 Strains and Primers

## Acknowledgements

We thank Xuewen Pan for the kind gift of Variomics libraries, and Guillermo Calero for one of the Pol II purifications used here. We thank Alex Francette and Karen Arndt for the gift of *S. pombe* chromatin and for advice and sharing of scripts for ChIP-seq analyses. We thank Allyson O’Donnell for advice on analysis of *YAP1-GFP* imaging and data analysis. We acknowledge funding from NIH R01GM097260 and R35GM144116 to CDK and also Welch Foundation Grant A-1763 to CDK for support on this project while our lab was at Texas A&M University. We acknowledge NSF for funding a Research Experience for Undergraduates site grant DBI-0851611 to Texas A&M Department of Biochemistry and Biophysics for support of AJL, SU, and EMP.

## Data availability

Sequencing data are available at the Sequence Read Archive under Bioproject. Scripts for data analysis are found at https://github.com/Kaplan-Lab-Pitt/Thiolutin.

---

**Figure S1.**
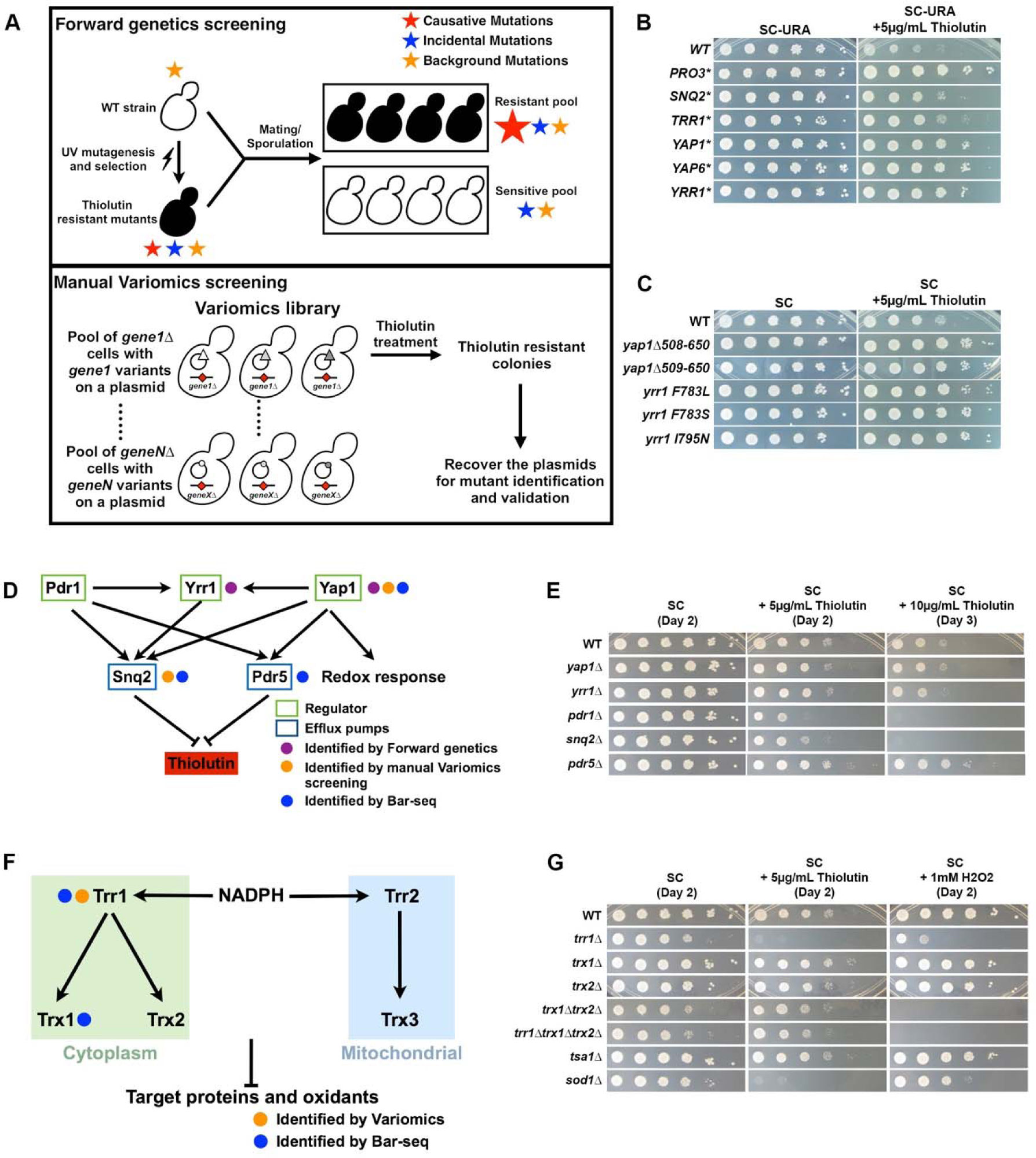
Mutations in multiple drug resistance (MDR) and oxidative stress responses (OSR) pathways alters cellular sensitivity to thiolutin. (A) Schematic diagram of the two independent genetic screens for thiolutin resistant mutants. Top: For the forward genetic screening, WT yeast strain (CKY457) was UV mutagenized and screened for thiolutin resistant mutants, and mutations were identified by whole-genome sequencing after bulk segregant analysis. Bottom: The gene-specific Variomics libraries consist of many mutants for a given gene in a yeast cell pool, and Variomics libraries for > 4000 genes were further pooled together for thiolutin resistance screening. Plasmids from the validated thiolutin resistant mutants were recovered for Sanger sequencing to identify both the gene and potential mutations. (B) Reproducibly-isolated thiolutin-resistant mutants from the forward genetics and manual Variomics screening. (C) Forward genetics isolated thiolutin-resistant alleles of *YAP1* and *YRR1*. (D) Schematic diagram summarizing a partial regulatory network of multiple drug resistance, with isolated resistant mutants indicated. The reported or hypothesized functional interactions are indicated by pointed (activation) or blunt-end (inhibition) arrows. (E) Distinct thiolutin sensitivity of MDR deficient mutants. *pdr1*Δ and *snq2*Δ are sensitive to thiolutin, whereas other tested MDR deficient strains retain resistance. (F) Schematic diagram summarizing the regulatory network of thioredoxin system in yeast, with isolated resistant mutants indicated. (G) Distinct thiolutin sensitivity of some OSR deficient mutants.

**Figure S2.**
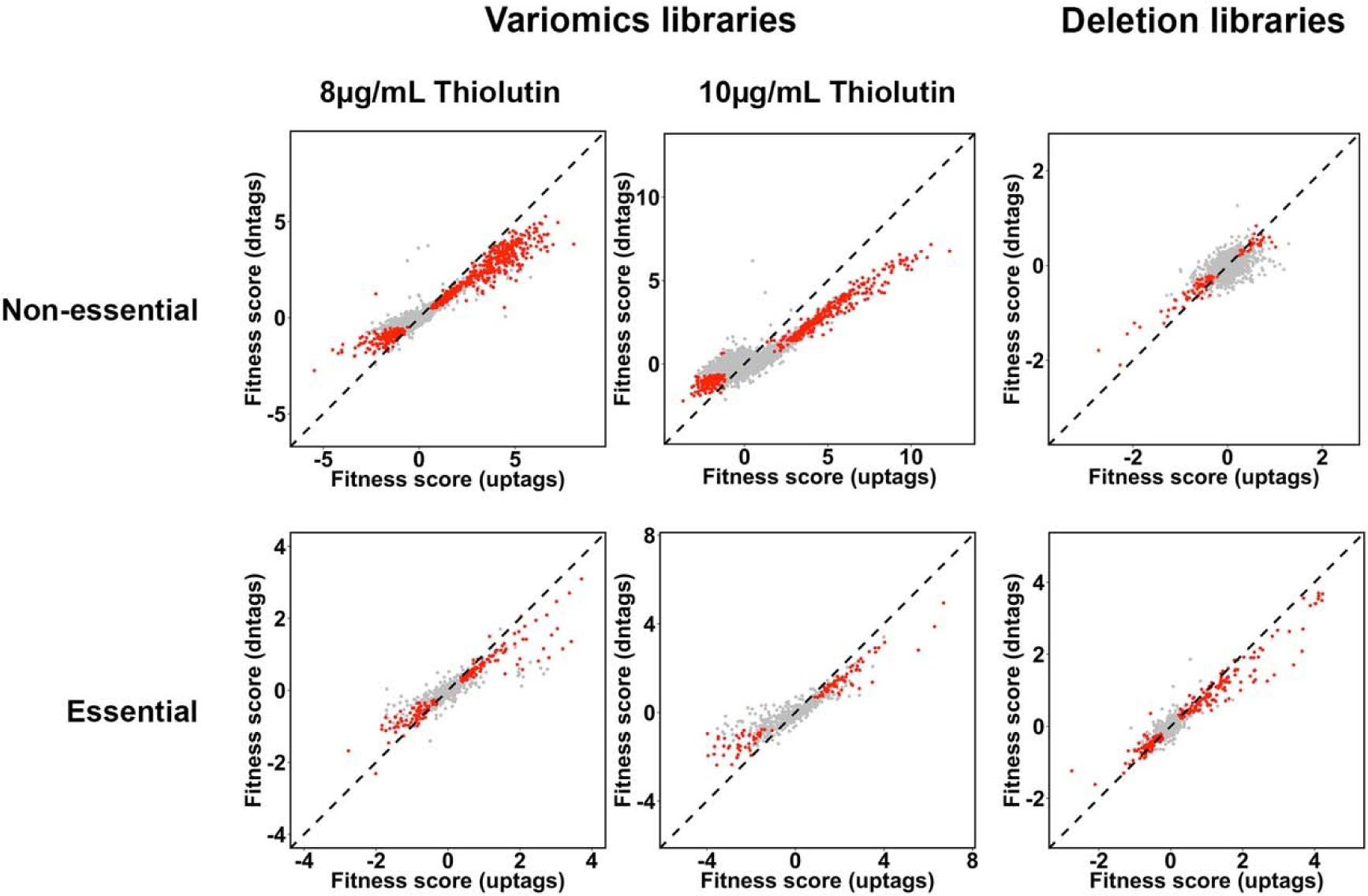
Correlation between uptag and downtags results in Bar-seq. Xyplots showing the fitness score (log2 of the fold change in relative abundance) computed from both uptag and downtags. Mutants are declared significantly resistant or sensitive in both tags are colored in red, whereas others are colored in light grey. y=x is shown as a dashed line.

**Figure S3.**
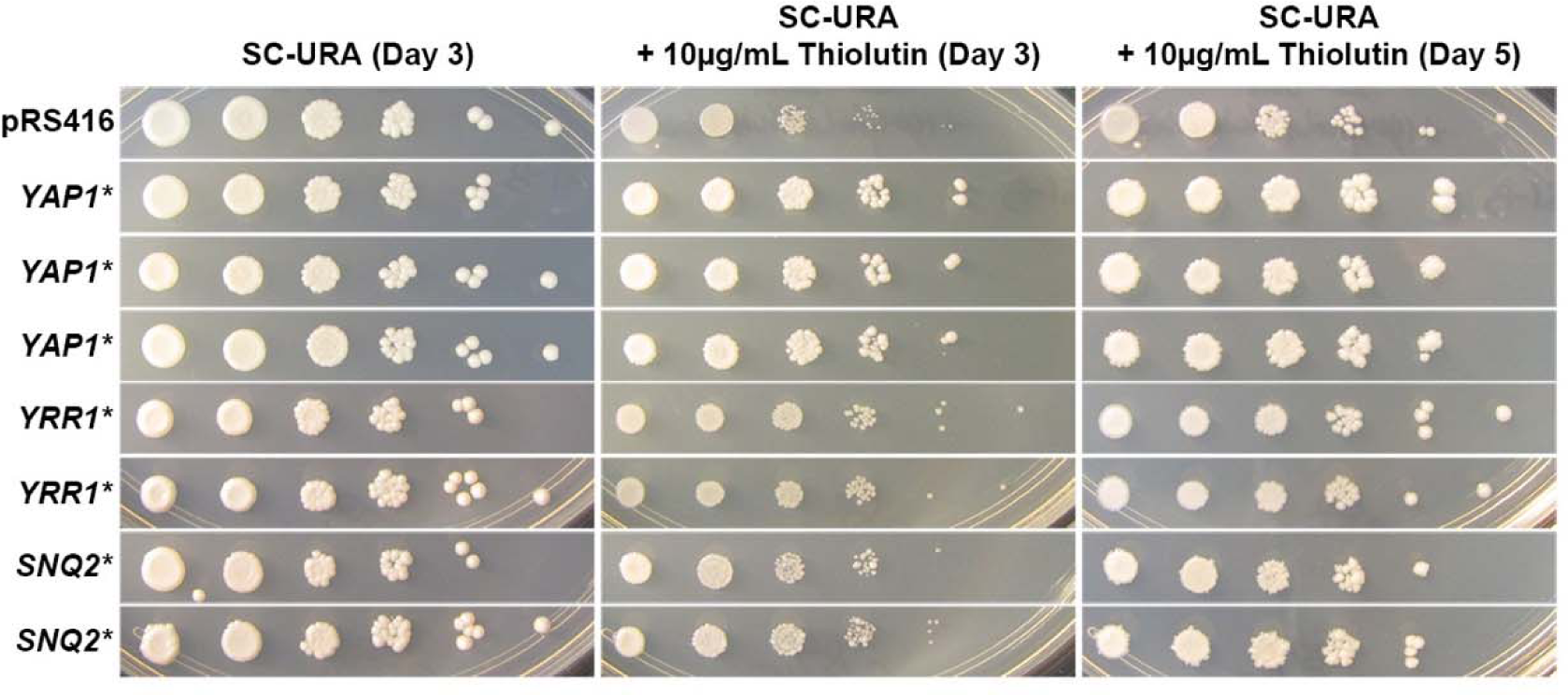
Isolated thiolutin resistant MDR Variomics candidates are dominant or dosage dependent. Plasmids recovered from the Variomics candidates are transformed into WT yeast strain along with an empty vector (pRS416) control, thiolutin resistance are assessed for at least two independent transformants.

**Figure S4.**
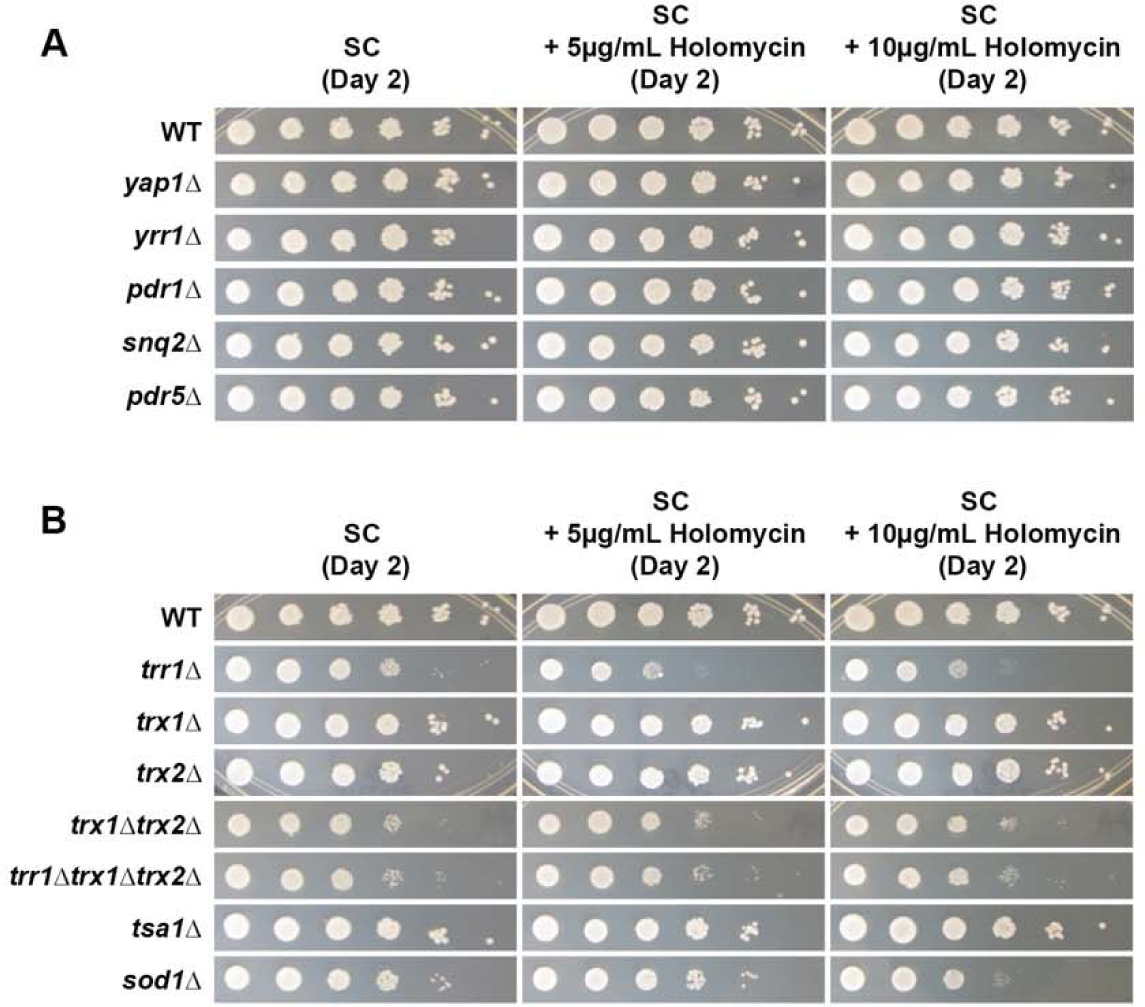
Tested MDR and OSR deficient mutants do not confer the same hypersensitivity to holomycin as to thiolutin. (A) Plate growth assays of serially diluted MDR deficient mutants on SC complete media with indicated holomycin concentration. (B) Plate growth assays of serially diluted OSR deficient mutants on SC complete media with indicated holomycin concentration.

**Figure S5.**
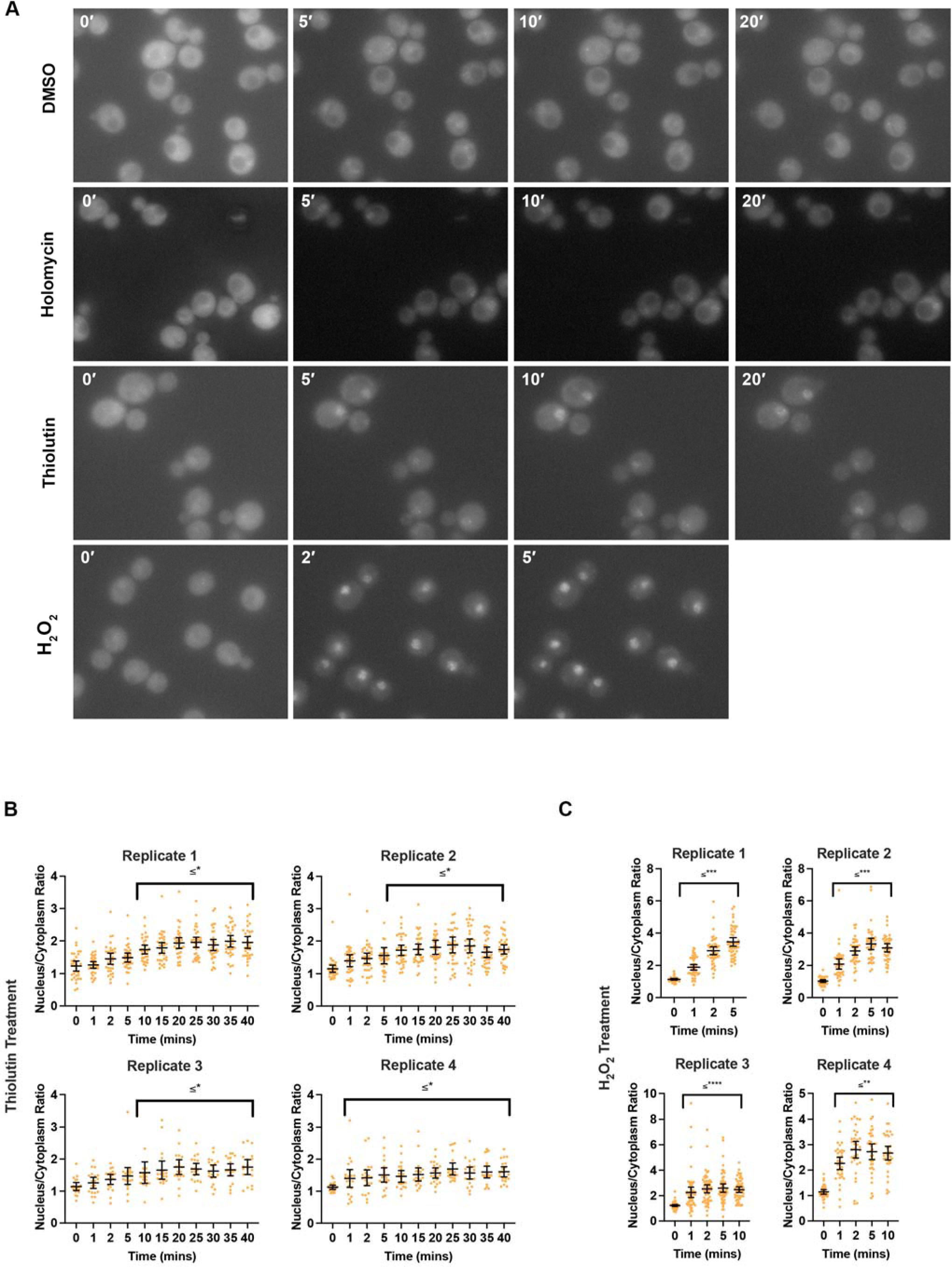
Thiolutin induces nuclear localization of the redox-sensitive Yap1 transcription activator. (A) Representative images of Yap1 relocalization to the nucleus after thiolutin or peroxide treatment. An EGFP-taggedYap1 strain was treated with 1% DMSO (control), 10µg/mL thiolutin, 10µg/mL holomycin, or 0.4mM H_2_O_2_. GFP fluorescence was monitored after treatment at the indicated time points. (B) Quantification of nuclear/cytoplasmic EGFP signal ratio for individual cells after 10µg/mL thiolutin treatment, for four independent biological replicates. Background was subtracted when nuclear and cytoplasmic signal were quantified. Error bars represent mean +/− the 95% confidence interval. (C) Quantification of nuclear/cytoplasmic EGFP signal ratio for individual cells after 0.4mM H_2_O_2_ treatment, in four different biological replicates. Quantification and error bars are as described in (B). Statistical test was Friedman’s paired test for each time point versus the 0 minute point for relevant cells (Dunn’s correction for multiple comparisons). Adjusted P-values indicated as *,**,***, **** were * ≤0.0332, ** ≤0.0021, *** ≤0.00002, **** <0.00001. “ ≤ *” indicates all P values were at * or below.

**Figure S6.**
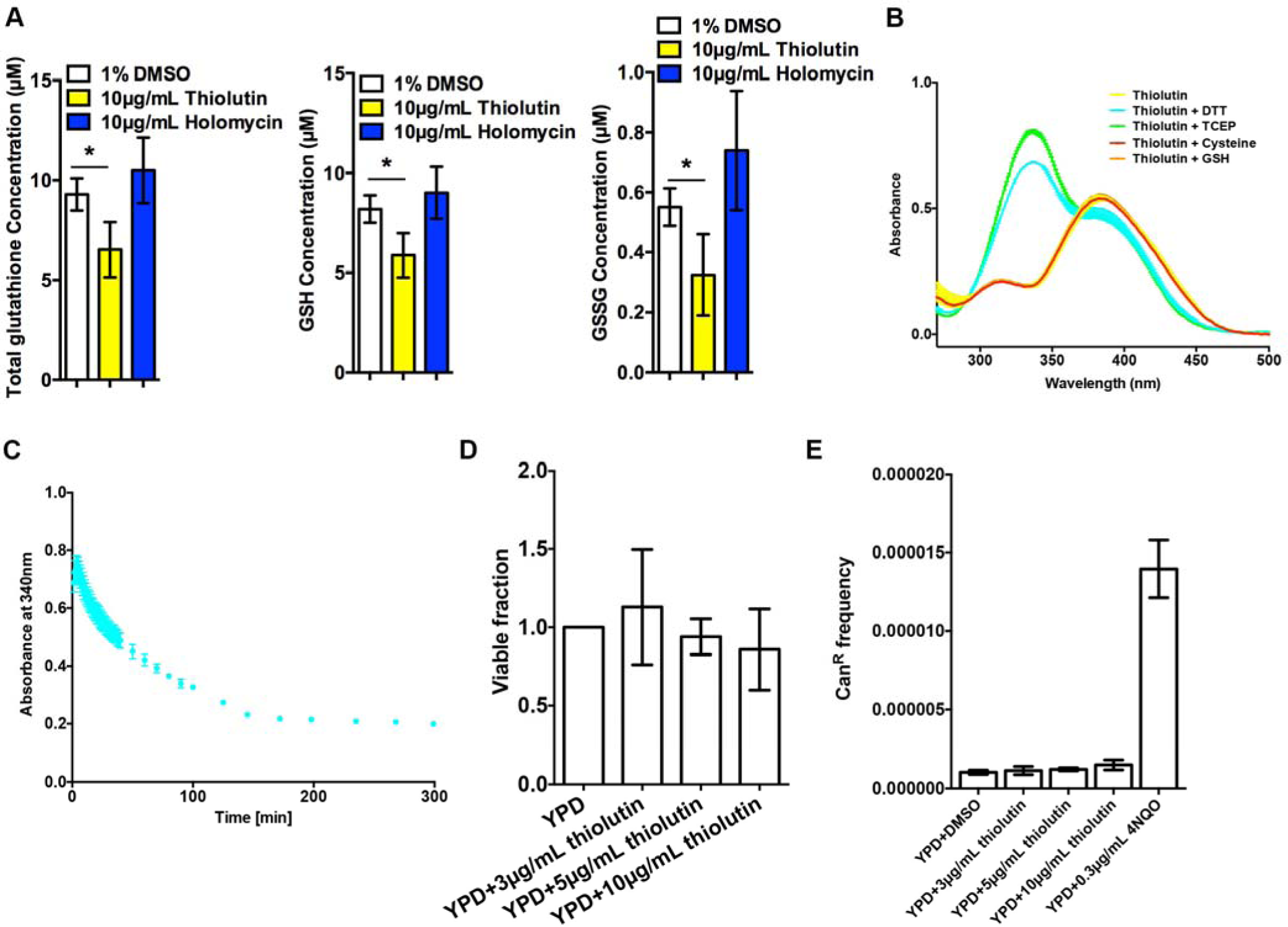
Thiolutin appears to induce oxidative stress partially through redox cycling. (A) Thiolutin depletes total glutathione *in vivo*. Growing WT yeast cultures are washed and treated with indicated conditions at 30°C for an hour. Reduced glutathione (GSH) and total glutathione are measured in three independent replicates, and the oxidized glutathione (GSSG) are calculated. The error bars represent standard deviation of the mean. *p≤0.01 (Two-tailed paired t-test). (B) Thiolutin can be reduced by DTT and TCEP, but not glutathione or cysteine *in vitro*. The reactions were performed in 100mM phosphate buffer (pH 6.5) with equivalent molar reductant with thiolutin. UV spectra were measured 1 minute after the reactions. Three independent replicates were performed, and the error bars represent standard deviation of the mean. (C) Reduced thiolutin is spontaneously re-oxidized when exposed to air. The reactions were performed as described in Figure 3D, and UV absorbance at 340nm plotted over a time course. Three independent replicates were performed and the error bars represent standard deviation of the mean. (D) Acute exposure to thiolutin does not affect yeast viability. (E) Thiolutin does not cause DNA damage, as measured by mutation frequency at *CAN1* gene.

**Figure S7.**
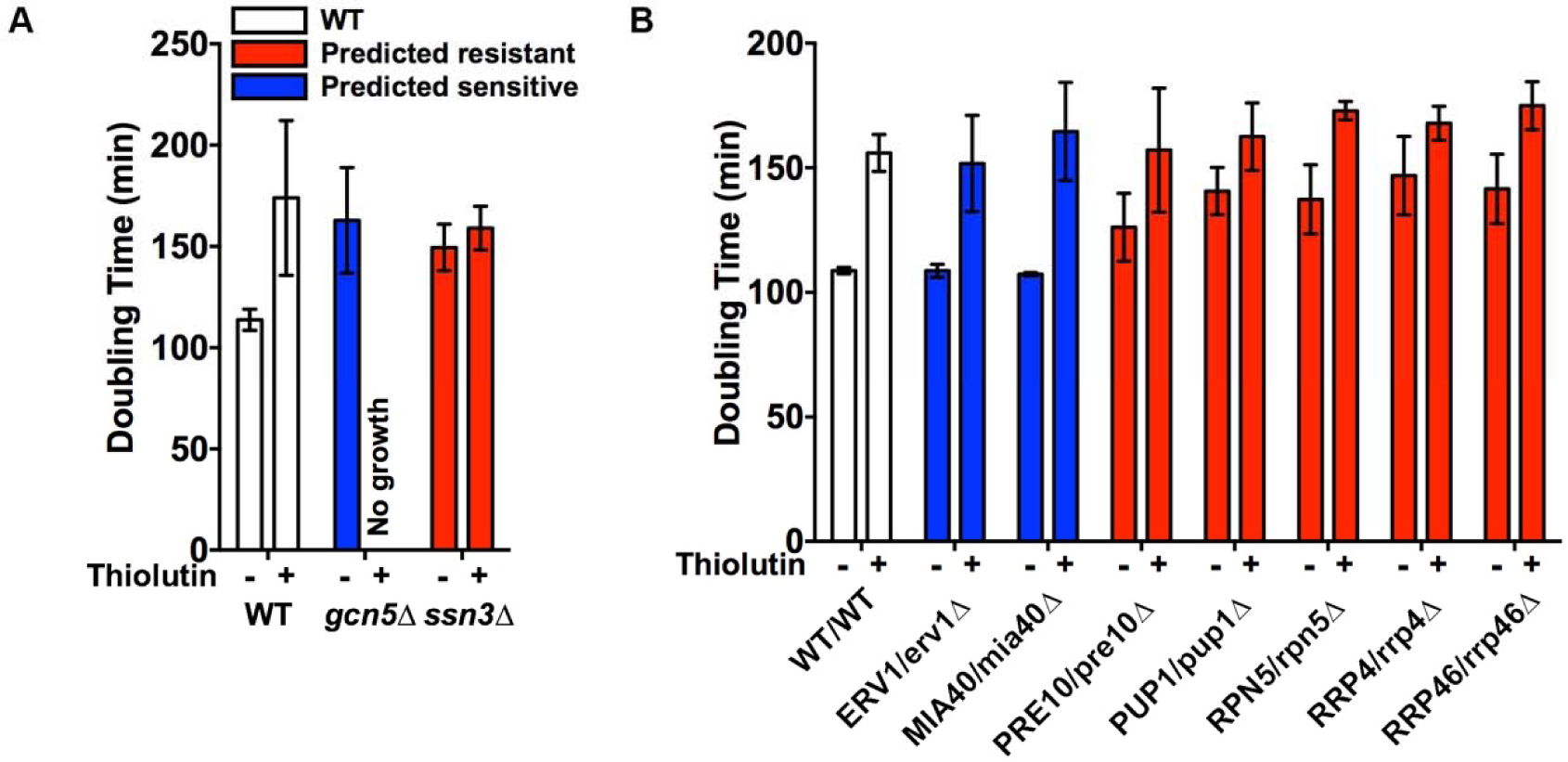
Validation of several statistically significantly resistant or sensitive mutants. Mutants were reconstructed in haploid BY4741 or diploid BY4743 backgrounds originally used to construct the deletion libraries, and mid-log growth at 30°C was assessed using a Tecan plate reader. Three independent repeats were performed and the error bars represent standard deviation of the mean. We failed to validate the observed slight sensitivity in *ERV1/erv1*Δ and *MIA40/mia40*Δ, two strains that were statistically significantly sensitive in Bar-seq but the sensitivity was very slight.

**Figure S8.**
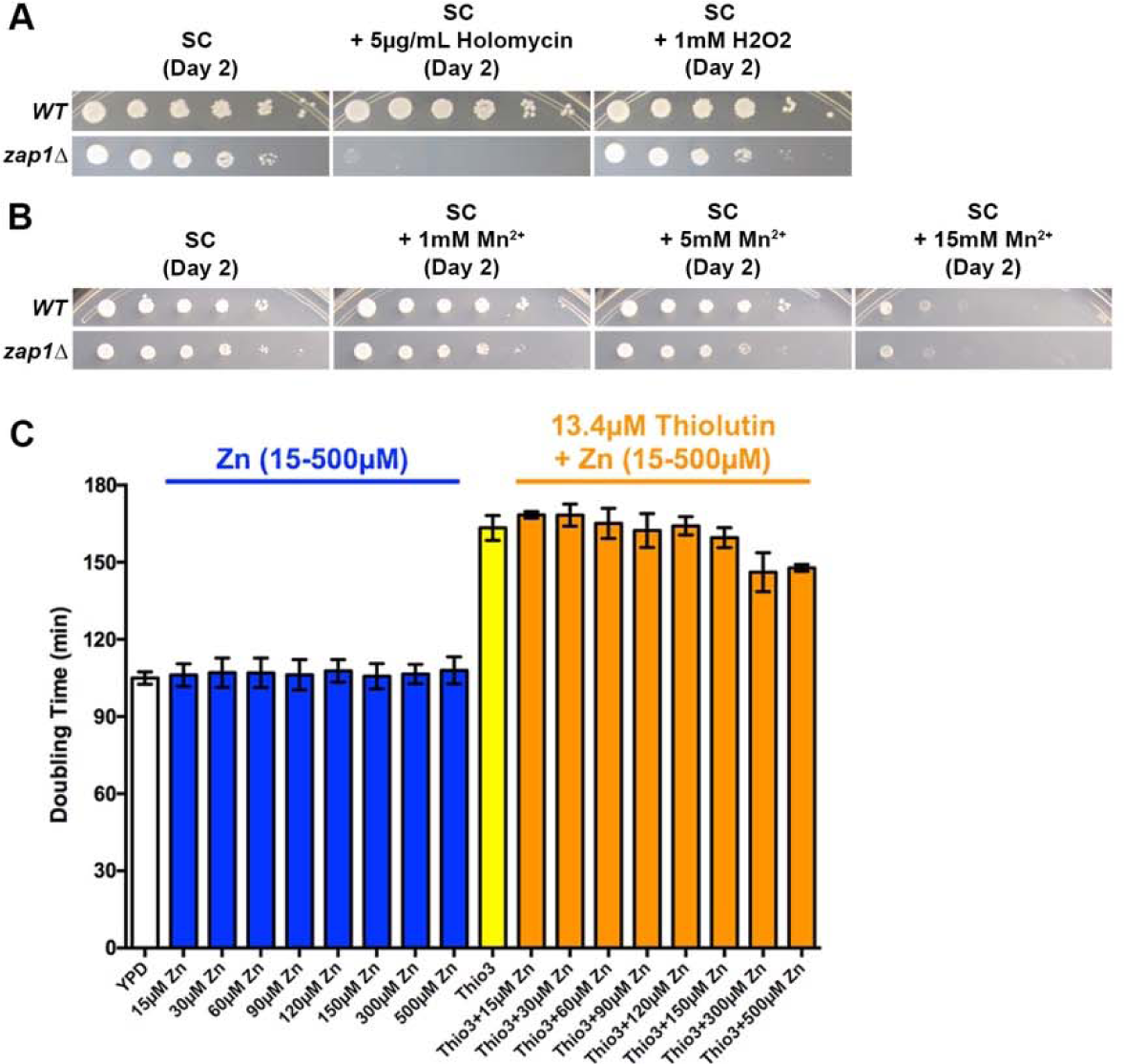
Modulation of intracellular Zn^2+^ alters holomycin and thiolutin sensitivity. (A) *zap1*Δ is hypersensitive to holomycin, but not H_2_O_2_. (B) *zap1*Δ doesn’t confer sensitivity to Mn^2+^. (C) Zn^2+^ supplementation partially suppresses thiolutin sensitivity. Doubling time were derived from growth curves measured by a Tecan plate reader. Three independent repeats were performed, and the error bars represent standard deviation of the mean.

**Figure S9.**
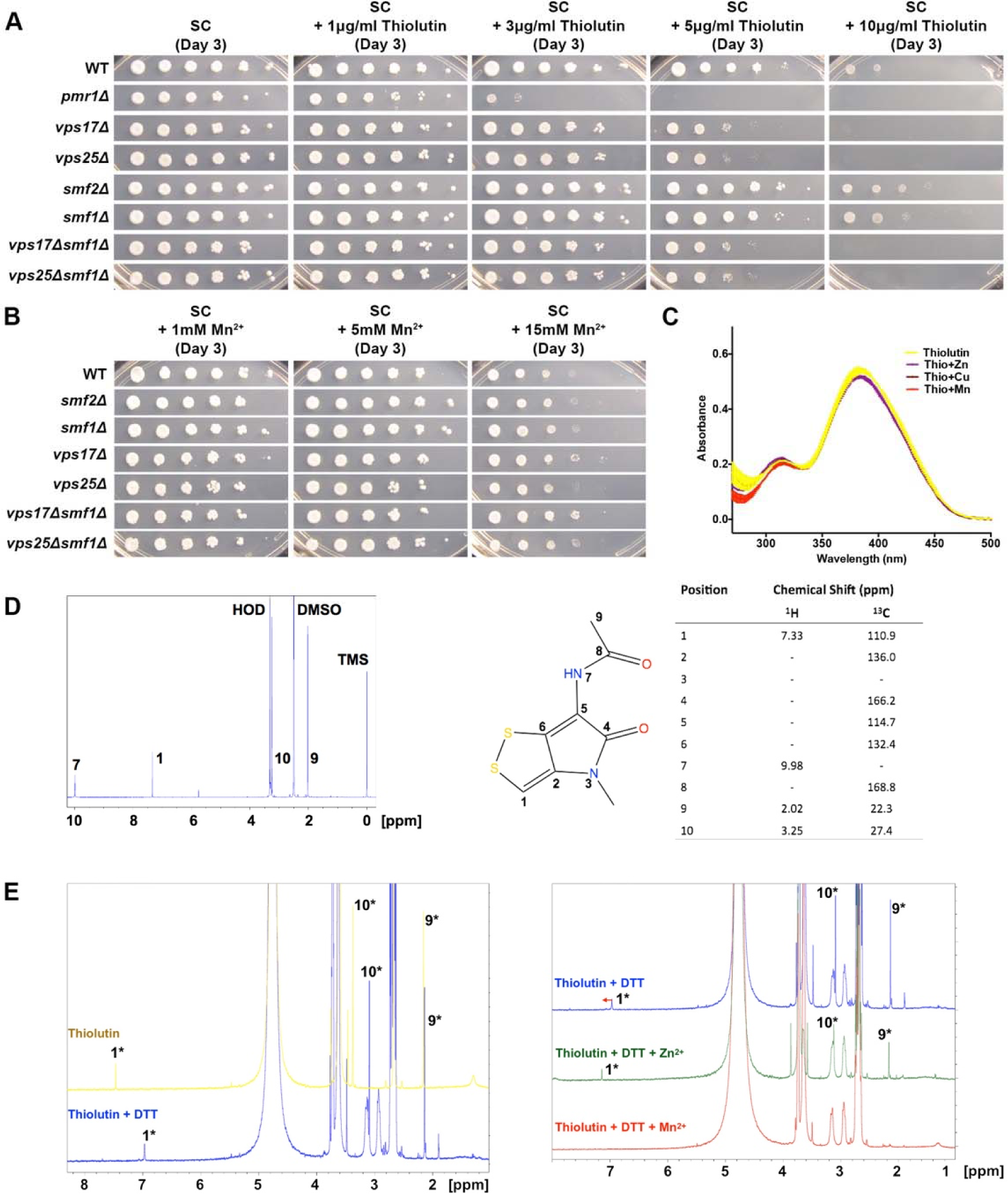
Mn^2+^ may modulate the sensitivity to reduced thiolutin *in vivo* and directly interact with thiolutin *in vitro*. (A) *pmr1*Δ is hyper-sensitive to thiolutin, while *vps17*Δ and *vps25*Δ appear to confer thiolutin sensitivity independent of Smf1-dependent Mn^2+^ trafficking. (B) *vps17*Δ, *vps25*Δ, *smf1*Δ and *smf2*Δ do not confer sensitivity to Mn^2+^ in our strain background and assays. (C) Zn^2+^, Cu^2+^ and Mn^2+^ do not alter the UV absorbance profile of thiolutin. (D) Assigned 1H-NMR spectrum of thiolutin. (E) Left: Change in thiolutin spectra immediately after reduction by DTT; Right: Changes in DTT-reduced thiolutin chemical shifts after addition of Zn^2+^ (diamagnetic) or Mn^2+^ (paramagnetic).

**Figure S10.**
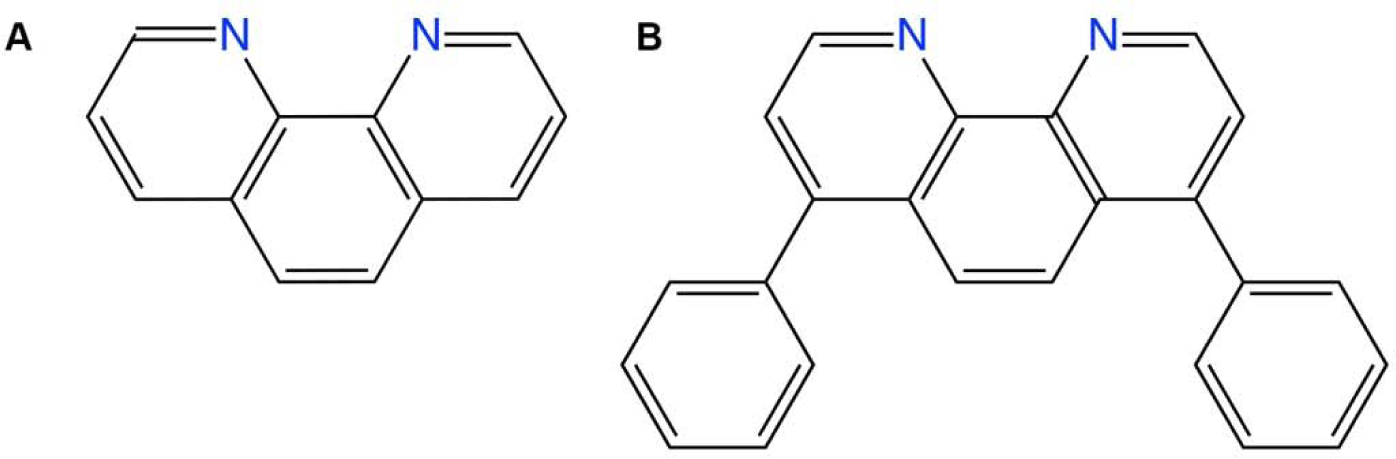
Structures of 1,10-phenanthroline and bathophenanthroline. (A) 1,10-phenanthroline. (B) bathophenanthroline

**Figure S11.**
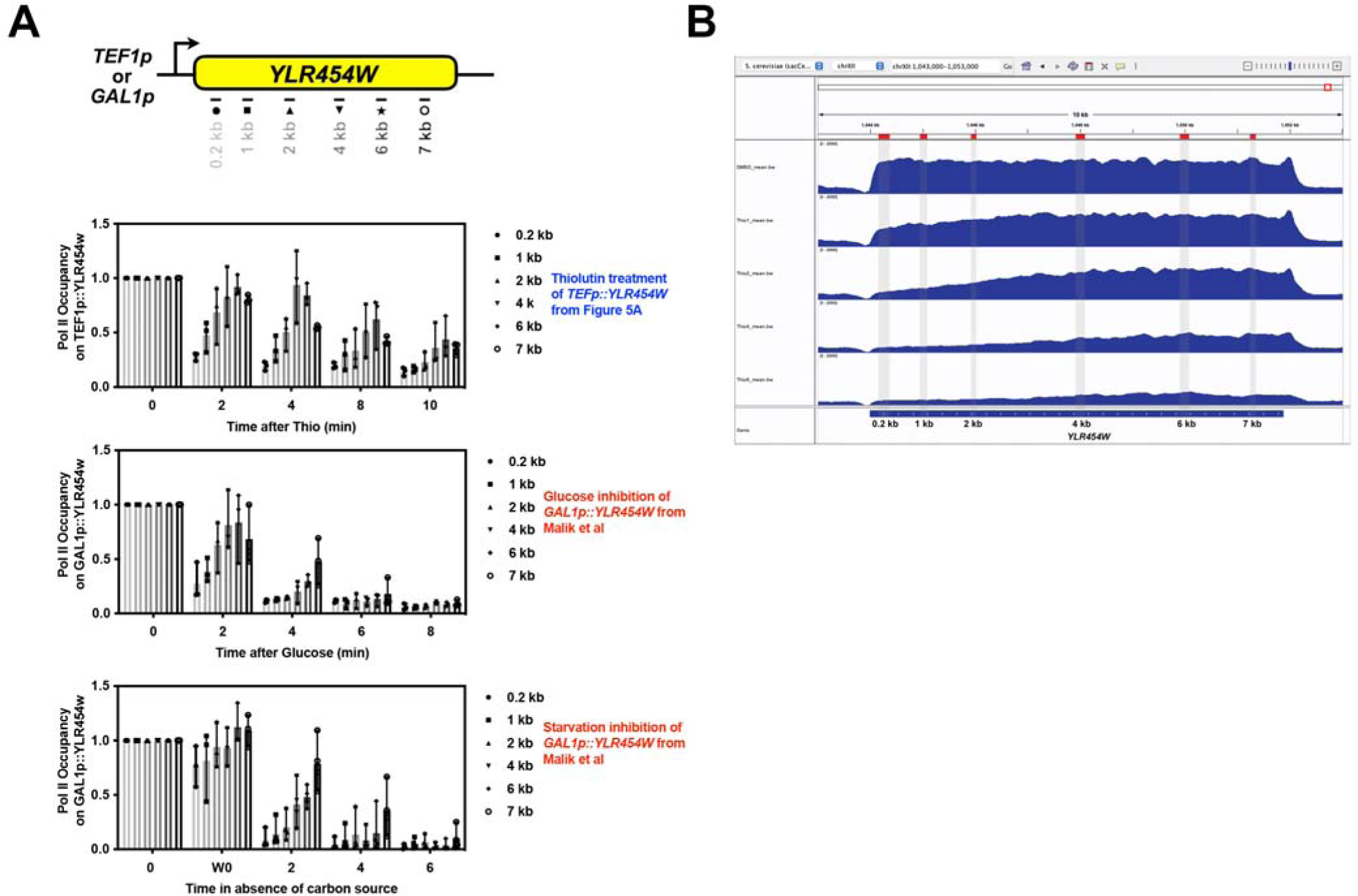
Comparison of Pol II occupancy on *YLR454W* after thiolutin treatment to other forms of transcription inhibition. (A) Pol II occupancy on *TEF1p::YLR454W* from Figure 5A (top) compared with Pol II occupancy changes upon glucose inhibition of *GAL1p::YLR454W* (middle) or carbon starvation (bottom) from Malik *et al* (ref. 39) as determined by ChIP-qPCR. Note that thiolutin treatment goes to ten minutes and Pol II has not cleared the 3′ end of the gene while inhibition by other methods shows complete clearance at or before six minutes. (B) Browser tracks showing ChIP-seq for Rpb3-FLAG (Figure 5) for *TEF1p::YLR454W* upon thiolutin treatment recapitulating ChIP-qPCR from (A). Gray highlights are positions of PCR amplicons used in (A) and in Malik *et al* (ref. 39).

**Figure S12.**
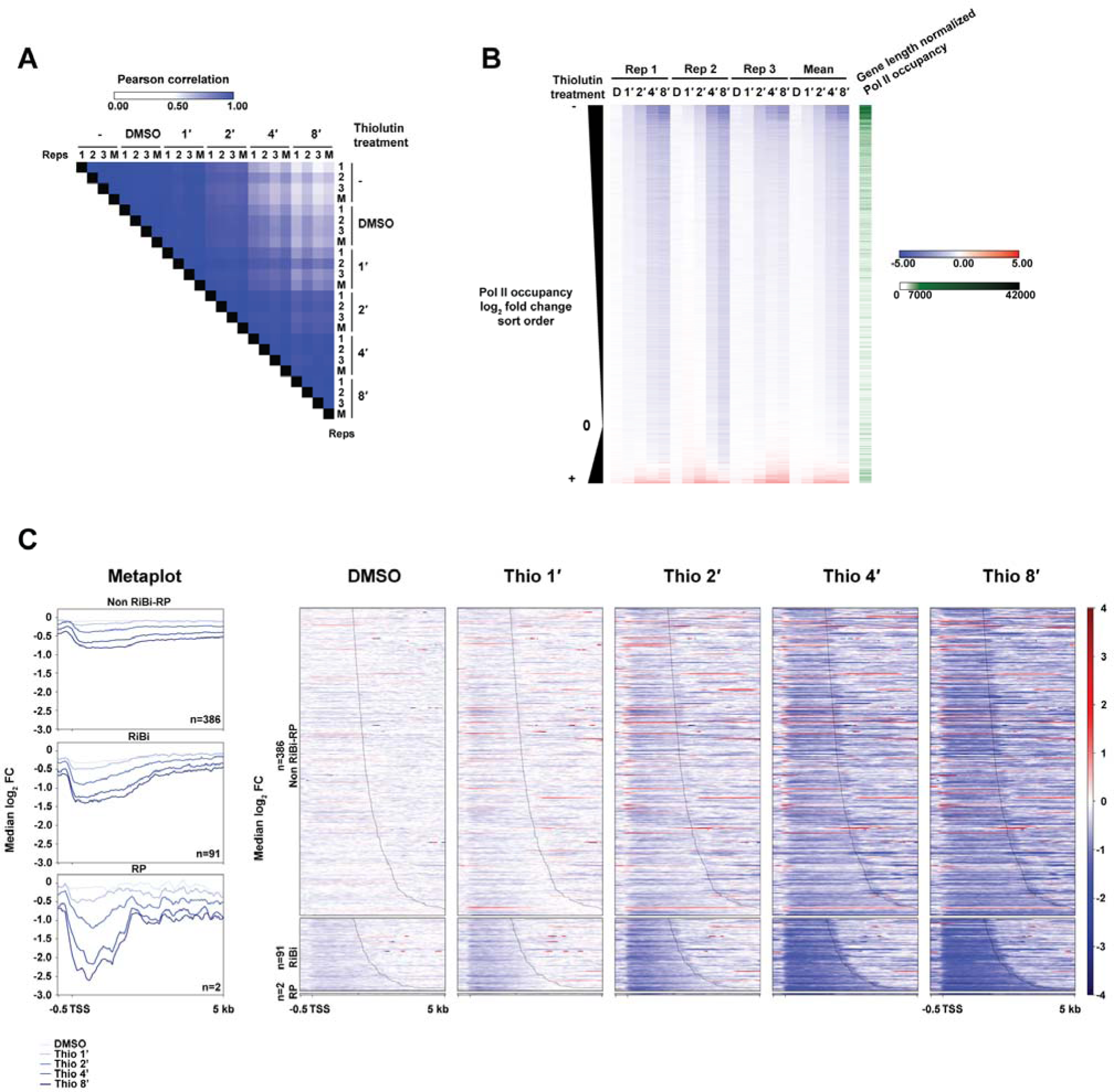
ChIP-seq for Pol II from thiolutin treated yeast. (A) Biological replicates of thiolutin treatment are reproducible. Heat map of Pearson r values for normalized Pol II occupancy/gene length for annotated yeast genes. Three biological replicates (“1”,”2”,”3”) and the mean of the replicates (“M”) for untreated cells (−), DMSO-treated control cells, and thiolutin treatment (10µg/mL) for 1, 2, 4, or 8 minutes are arranged in a heat map. Pearson r values are indicated by the color scale for comparison of samples on the x and y axes. (B) Yeast genes rank ordered from greatest decrease in Pol II occupancy to greatest increase for biological replicates with log_2_ fold change in spike-in normalized Pol II occupancy/gene length shown in a heat map. (C) Yeast genes from the top 40% based on Pol II occupancy in untreated cells that are ≥1.5 kb in length were divided into non-RP/RiBi genes (n=386), RiBi genes (n=91), and RP genes (n=2) and ordered by length (Right panels). Median log_2_ fold change in Pol II occupancy between treatment (DMSO or thiolutin) and untreated conditions is shown. Left panels are the metaplots for change in Pol II occupancy over time of thiolutin treatment.

## References

1. Li, B. and Walsh, C.T. (2011) Streptomyces clavuligerus HlmI is an intramolecular disulfide-forming dithiol oxidase in holomycin biosynthesis. Biochemistry, 50, 4615–4622.

2. Li, B., Wever, W.J., Walsh, C.T. and Bowers, A.A. (2014) Dithiolopyrrolones: biosynthesis, synthesis, and activity of a unique class of disulfide-containing antibiotics. Natural product reports, 31, 905–923.

3. Rossini, C., Taylor, W., Fagan, T. and Hastings, J.W. (2003) Lifetimes of mRNAs for clock-regulated proteins in a dinoflagellate. Chronobiol Int, 20, 963–976.

4. Grigull, J., Mnaimneh, S., Pootoolal, J., Robinson, M.D. and Hughes, T.R. (2004) Genome-wide analysis of mRNA stability using transcription inhibitors and microarrays reveals posttranscriptional control of ribosome biogenesis factors. Molecular and cellular biology, 24, 5534–5547.

5. Kebaara, B.W., Nielsen, L.E., Nickerson, K.W. and Atkin, A.L. (2006) Determination of mRNA half-lives in Candida albicans using thiolutin as a transcription inhibitor. Genome, 49, 894–899.

6. Morey, J.S. and Van Dolah, F.M. (2013) Global analysis of mRNA half-lives and de novo transcription in a dinoflagellate, Karenia brevis. PLoS One, 8, e66347.

7. Bergmann, R. (1989) Thiolutin inhibits utilization of glucose and other carbon sources in cells of Escherichia coli. Antonie Van Leeuwenhoek, 55, 143–152.

8. Eshleman, N., Luo, X., Capaldi, A. and Buchan, J.R. (2020) Alterations of signaling pathways in response to chemical perturbations used to measure mRNA decay rates in yeast. Rna, 26, 10–18.

9. Pelechano, V. and Perez-Ortin, J.E. (2008) The transcriptional inhibitor thiolutin blocks mRNA degradation in yeast. Yeast, 25, 85–92.

10. Monje-Casas, F., Michan, C. and Pueyo, C. (2004) Absolute transcript levels of thioredoxin- and glutathione-dependent redox systems in Saccharomyces cerevisiae: response to stress and modulation with growth. Biochem J, 383, 139–147.

11. Lauinger, L., Li, J., Shostak, A., Cemel, I.A., Ha, N., Zhang, Y., Merkl, P.E., Obermeyer, S., Stankovic-Valentin, N. and Schafmeier, T. (2017) Thiolutin is a zinc chelator that inhibits the Rpn11 and other JAMM metalloproteases. Nature Chemical Biology, 13, 709–714.

12. Chan, A.N., Shiver, A.L., Wever, W.J., Razvi, S.Z., Traxler, M.F. and Li, B. (2017) Role for dithiolopyrrolones in disrupting bacterial metal homeostasis. Proceedings of the National Academy of Sciences of the United States of America, 114, 2717–2722.

13. Fuchs, A.C.D., Maldoner, L., Wojtynek, M., Hartmann, M.D. and Martin, J. (2018) Rpn11-mediated ubiquitin processing in an ancestral archaeal ubiquitination system. Nat Commun, 9, 2696.

14. Baell, J.B. (2011) Redox-active nuisance screening compounds and their classification. Drug Discovery Today, 16, 840–841.

15. Johnston, P.A. (2011) Redox cycling compounds generate H2O2 in HTS buffers containing strong reducing reagents--real hits or promiscuous artifacts? Curr Opin Chem Biol, 15, 174–182.

16. Albini, F., Bormann, S., Gerschel, P., Ludwig, V.A. and Neumann, W. (2023) Dithiolopyrrolones are Prochelators that are Activated by Glutathione. Chemistry–A European Journal, 29, e202202567.

17. Khachatourians, G.G. and Tipper, D.J. (1974) In vivo effect of thiolutin on cell growth and macromolecular synthesis in Escherichia coli. Antimicrob Agents Chemother, 6, 304–310.

18. Joshi, A., Verma, M. and Chakravorty, M. (1982) Thiolutin-resistant mutants of Salmonella typhimurium. Antimicrob Agents Chemother, 22, 541–547.

19. Roza, J., Blanco, M.G., Hardisson, C. and Salas, J.A. (1986) Self-resistance in actinomycetes producing inhibitors of RNA polymerase. The Journal of antibiotics, 39, 609–612.

20. Tipper, D.J. (1973) Inhibition of yeast ribonucleic acid polymerases by thiolutin. J Bacteriol, 116, 245–256.

21. Sivasubramanian, N. and Jayaraman, R. (1976) Thiolutin resistant mutants of Escherichia coli are they RNA chain initiation mutants? Molecular and General Genetics MGG, 145, 89–96.

22. Oliva, B., O’Neill, A., Wilson, J.M., O’Hanlon, P.J. and Chopra, I. (2001) Antimicrobial properties and mode of action of the pyrrothine holomycin. Antimicrob Agents Chemother, 45, 532–539.

23. Mukhopadhyay, J., Das, K., Ismail, S., Koppstein, D., Jang, M., Hudson, B., Sarafianos, S., Tuske, S., Patel, J. and Jansen, R. (2008) The RNA polymerase “switch region” is a target for inhibitors. Cell, 135, 295–307.

24. Wang, D. (2008) Ensemble fluorescence resonance energy transfer analysis of RNA polymerase clamp conformation. Rutgers The State University of New Jersey-New Brunswick.

25. Smith, A.M., Heisler, L.E., Mellor, J., Kaper, F., Thompson, M.J., Chee, M., Roth, F.P., Giaever, G. and Nislow, C. (2009) Quantitative phenotyping via deep barcode sequencing. Genome Res, 19, 1836–1842.

26. Lee, A.Y., St Onge, R.P., Proctor, M.J., Wallace, I.M., Nile, A.H., Spagnuolo, P.A., Jitkova, Y., Gronda, M., Wu, Y., Kim, M.K. et al. (2014) Mapping the cellular response to small molecules using chemogenomic fitness signatures. Science, 344, 208–211.

27. Parsons, A.B., Lopez, A., Givoni, I.E., Williams, D.E., Gray, C.A., Porter, J., Chua, G., Sopko, R., Brost, R.L., Ho, C.H. et al. (2006) Exploring the mode-of-action of bioactive compounds by chemical-genetic profiling in yeast. Cell, 126, 611–625.

28. Perrin, D.M., Pearson, L., Mazumder, A. and Sigman, D.S. (1994) Inhibition of prokaryotic and eukaryotic transcription by the 2:1 2,9-dimethyl-1,10-phenanthroline-cuprous complex, a ligand specific for open complexes. Gene, 149, 173–178.

29. Rohde, J.R., Bastidas, R., Puria, R. and Cardenas, M.E. (2008) Nutritional control via Tor signaling in Saccharomyces cerevisiae. Current opinion in microbiology, 11, 153–160.

30. Shore, D., Zencir, S. and Albert, B. (2021) Transcriptional control of ribosome biogenesis in yeast: links to growth and stress signals. Biochemical Society transactions, 49, 1589–1599.

31. Janke, C., Magiera, M.M., Rathfelder, N., Taxis, C., Reber, S., Maekawa, H., Moreno-Borchart, A., Doenges, G., Schwob, E., Schiebel, E. et al. (2004) A versatile toolbox for PCR-based tagging of yeast genes: new fluorescent proteins, more markers and promoter substitution cassettes. Yeast, 21, 947–962.

32. Chee, M.K. and Haase, S.B. (2012) New and redesigned pRS plasmid shuttle vectors for genetic manipulation of Saccharomyces cerevisiae. G3: Genes, Genomes, Genetics, 2, 515–526.

33. Huang, Z., Chen, K., Zhang, J., Li, Y., Wang, H., Cui, D., Tang, J., Liu, Y., Shi, X., Li, W. et al. (2013) A functional variomics tool for discovering drug-resistance genes and drug targets. Cell reports, 3, 577–585.

34. Langmead, B. and Salzberg, S.L. (2012) Fast gapped-read alignment with Bowtie 2. Nat Methods, 9, 357–359.

35. Li, H., Handsaker, B., Wysoker, A., Fennell, T., Ruan, J., Homer, N., Marth, G., Abecasis, G., Durbin, R. and Genome Project Data Processing, S. (2009) The Sequence Alignment/Map format and SAMtools. Bioinformatics, 25, 2078–2079.

36. Robinson, D.G., Chen, W., Storey, J.D. and Gresham, D. (2014) Design and analysis of Bar-seq experiments. G3 (Bethesda, Md.), 4, 11–18.

37. Robinson, M.D., McCarthy, D.J. and Smyth, G.K. (2010) edgeR: a Bioconductor package for differential expression analysis of digital gene expression data. bioinformatics, 26, 139–140.

38. de Hoon, M.J., Imoto, S., Nolan, J. and Miyano, S. (2004) Open source clustering software. Bioinformatics, 20, 1453–1454.

39. Malik, I., Qiu, C., Snavely, T. and Kaplan, C.D. (2017) Wide-ranging and unexpected consequences of altered Pol II catalytic activity in vivo. Nucleic Acids Res, 45, 4431–4451.

40. Kaplan, C.D., Jin, H., Zhang, I.L. and Belyanin, A. (2012) Dissection of Pol II trigger loop function and Pol II activity-dependent control of start site selection in vivo. PLoS genetics, 8, e1002627.

41. Qiu, C., Erinne, O.C., Dave, J.M., Cui, P., Jin, H., Muthukrishnan, N., Tang, L.K., Babu, S.G., Lam, K.C., Vandeventer, P.J. et al. (2016) High-Resolution Phenotypic Landscape of the RNA Polymerase II Trigger Loop. PLoS genetics, 12, e1006321.

42. Kaplan, C.D., Larsson, K.M. and Kornberg, R.D. (2008) The RNA polymerase II trigger loop functions in substrate selection and is directly targeted by alpha-amanitin. Mol Cell, 30, 547–556.

43. Bolger, A.M., Lohse, M. and Usadel, B. (2014) Trimmomatic: a flexible trimmer for Illumina sequence data. Bioinformatics, 30, 2114–2120.

44. Jeronimo, C., Poitras, C. and Robert, F. (2019) Histone Recycling by FACT and Spt6 during Transcription Prevents the Scrambling of Histone Modifications. Cell reports, 28, 1206–1218.e1208.

45. Zerbino, D.R., Johnson, N., Juettemann, T., Wilder, S.P. and Flicek, P. (2014) WiggleTools: parallel processing of large collections of genome-wide datasets for visualization and statistical analysis. Bioinformatics, 30, 1008–1009.

46. Ramírez, F., Ryan, D.P., Grüning, B., Bhardwaj, V., Kilpert, F., Richter, A.S., Heyne, S., Dündar, F. and Manke, T. (2016) deepTools2: a next generation web server for deep-sequencing data analysis. Nucleic acids research, 44, W160.

47. Kent, W.J., Zweig, A.S., Barber, G., Hinrichs, A.S. and Karolchik, D. (2010) BigWig and BigBed: enabling browsing of large distributed datasets. Bioinformatics, 26, 2204–2207.

48. Birkeland, S.R., Jin, N., Ozdemir, A.C., Lyons, R.H., Jr., Weisman, L.S. and Wilson, T.E. (2010) Discovery of mutations in Saccharomyces cerevisiae by pooled linkage analysis and whole-genome sequencing. Genetics, 186, 1127–1137.

49. Karim, A.S., Curran, K.A. and Alper, H.S. (2013) Characterization of plasmid burden and copy number in Saccharomyces cerevisiae for optimization of metabolic engineering applications. FEMS Yeast Res, 13, 107–116.

50. Kuge, S., Jones, N. and Nomoto, A. (1997) Regulation of yAP-1 nuclear localization in response to oxidative stress. EMBO J, 16, 1710–1720.

51. Wemmie, J.A., Steggerda, S.M. and Moye-Rowley, W.S. (1997) The Saccharomyces cerevisiae AP-1 protein discriminates between oxidative stress elicited by the oxidants H2O2 and diamide. Journal of Biological Chemistry, 272, 7908–7914.

52. Coleman, S.T., Epping, E.A., Steggerda, S.M. and Moye-Rowley, W.S. (1999) Yap1p activates gene transcription in an oxidant-specific fashion. Mol Cell Biol, 19, 8302–8313.

53. Kuge, S., Arita, M., Murayama, A., Maeta, K., Izawa, S., Inoue, Y. and Nomoto, A. (2001) Regulation of the yeast Yap1p nuclear export signal is mediated by redox signal-induced reversible disulfide bond formation. Molecular and cellular biology, 21, 6139–6150.

54. Gulshan, K., Rovinsky, S.A., Coleman, S.T. and Moye-Rowley, W.S. (2005) Oxidant-specific folding of Yap1p regulates both transcriptional activation and nuclear localization. Journal of Biological Chemistry, 280, 40524–40533.

55. Alarco, A.M., Balan, I., Talibi, D., Mainville, N. and Raymond, M. (1997) AP1-mediated multidrug resistance in Saccharomyces cerevisiae requires FLR1 encoding a transporter of the major facilitator superfamily. The Journal of biological chemistry, 272, 19304–19313.

56. Coleman, S.T., Tseng, E. and Moye-Rowley, W.S. (1997) Saccharomyces cerevisiae basic region-leucine zipper protein regulatory networks converge at the ATR1 structural gene. Journal of Biological Chemistry, 272, 23224–23230.

57. Gulshan, K. and Moye-Rowley, W.S. (2007) Multidrug resistance in fungi. Eukaryot Cell, 6, 1933–1942.

58. Meyer, Y., Buchanan, B.B., Vignols, F. and Reichheld, J.P. (2009) Thioredoxins and glutaredoxins: unifying elements in redox biology. Annu Rev Genet, 43, 335–367.

59. Winzeler, E.A., Shoemaker, D.D., Astromoff, A., Liang, H., Anderson, K., Andre, B., Bangham, R., Benito, R., Boeke, J.D., Bussey, H. et al. (1999) Functional characterization of the S. cerevisiae genome by gene deletion and parallel analysis. Science, 285, 901–906.

60. Machado, A.K., Morgan, B.A. and Merrill, G.F. (1997) Thioredoxin reductase-dependent inhibition of MCB cell cycle box activity in Saccharomyces cerevisiae. The Journal of biological chemistry, 272, 17045–17054.

61. Carmel-Harel, O., Stearman, R., Gasch, A.P., Botstein, D., Brown, P.O. and Storz, G. (2001) Role of thioredoxin reductase in the Yap1p-dependent response to oxidative stress in Saccharomyces cerevisiae. Mol Microbiol, 39, 595–605.

62. Trotter, E.W. and Grant, C.M. (2002) Thioredoxins are required for protection against a reductive stress in the yeast Saccharomyces cerevisiae. Mol Microbiol, 46, 869–878.

63. Hacioglu, E., Esmer, I., Fomenko, D.E., Gladyshev, V.N. and Koc, A. (2010) The roles of thiol oxidoreductases in yeast replicative aging. Mech Ageing Dev, 131, 692–699.

64. Ragu, S., Dardalhon, M., Sharma, S., Iraqui, I., Buhagiar-Labarchede, G., Grondin, V., Kienda, G., Vernis, L., Chanet, R., Kolodner, R.D. et al. (2014) Loss of the thioredoxin reductase Trr1 suppresses the genomic instability of peroxiredoxin tsa1 mutants. PLoS One, 9, e108123.

65. Grant, C.M., Perrone, G. and Dawes, I.W. (1998) Glutathione and Catalase Provide Overlapping Defenses for Protection against Hydrogen Peroxide in the YeastSaccharomyces cerevisiae. Biochemical and biophysical research communications, 253, 893–898.

66. Castro, F.A., Mariani, D., Panek, A.D., Eleutherio, E.C. and Pereira, M.D. (2008) Cytotoxicity mechanism of two naphthoquinones (menadione and plumbagin) in Saccharomyces cerevisiae. PLoS One, 3, e3999.

67. Toledano, M.B., Delaunay-Moisan, A., Outten, C.E. and Igbaria, A. (2013) Functions and cellular compartmentation of the thioredoxin and glutathione pathways in yeast. Antioxid Redox Signal, 18, 1699–1711.

68. Jung, U., Zheng, X., Yoon, S.O. and Chung, A.S. (2001) Se-methylselenocysteine induces apoptosis mediated by reactive oxygen species in HL-60 cells. Free Radic Biol Med, 31, 479–489.

69. Ravi, D. and Das, K.C. (2004) Redox-cycling of anthracyclines by thioredoxin system: increased superoxide generation and DNA damage. Cancer Chemother Pharmacol, 54, 449–458.

70. Jimenez, A., Tipper, D.J. and Davies, J. (1973) Mode of action of thiolutin, an inhibitor of macromolecular synthesis in Saccharomyces cerevisiae. Antimicrob Agents Chemother, 3, 729–738.

71. Dudley, A.M., Janse, D.M., Tanay, A., Shamir, R. and Church, G.M. (2005) A global view of pleiotropy and phenotypically derived gene function in yeast. Mol Syst Biol, 1, 2005 0001.

72. Pagani, M.A., Casamayor, A., Serrano, R., Atrian, S. and Arino, J. (2007) Disruption of iron homeostasis in Saccharomyces cerevisiae by high zinc levels: a genome-wide study. Mol Microbiol, 65, 521–537.

73. Ruotolo, R., Marchini, G. and Ottonello, S. (2008) Membrane transporters and protein traffic networks differentially affecting metal tolerance: a genomic phenotyping study in yeast. Genome biology, 9, R67.

74. Bleackley, M.R., Young, B.P., Loewen, C.J. and MacGillivray, R.T. (2011) High density array screening to identify the genetic requirements for transition metal tolerance in Saccharomyces cerevisiae. Metallomics, 3, 195–205.

75. Jiang, L., Cao, C., Zhang, L., Lin, W., Xia, J., Xu, H. and Zhang, Y. (2014) Cadmium-induced activation of high osmolarity glycerol pathway through its Sln1 branch is dependent on the MAP kinase kinase kinase Ssk2, but not its paralog Ssk22, in budding yeast. FEMS Yeast Res, 14, 1263–1272.

76. Acosta-Alvear, D., Cho, M.Y., Wild, T., Buchholz, T.J., Lerner, A.G., Simakova, O., Hahn, J., Korde, N., Landgren, O., Maric, I. et al. (2015) Paradoxical resistance of multiple myeloma to proteasome inhibitors by decreased levels of 19S proteasomal subunits. Elife, 4, e08153.

77. Tsvetkov, P., Mendillo, M.L., Zhao, J., Carette, J.E., Merrill, P.H., Cikes, D., Varadarajan, M., van Diemen, F.R., Penninger, J.M. and Goldberg, A.L. (2015) Compromising the 19S proteasome complex protects cells from reduced flux through the proteasome. Elife, 4, e08467.

78. Tsvetkov, P., Sokol, E., Jin, D., Brune, Z., Thiru, P., Ghandi, M., Garraway, L.A., Gupta, P.B., Santagata, S., Whitesell, L. et al. (2017) Suppression of 19S proteasome subunits marks emergence of an altered cell state in diverse cancers. Proceedings of the National Academy of Sciences of the United States of America, 114, 382–387.

79. Bandara, P.D., Flattery-O’Brien, J.A., Grant, C.M. and Dawes, I.W. (1998) Involvement of the Saccharomyces cerevisiae UTH1 gene in the oxidative-stress response. Curr Genet, 34, 259–268.

80. Wu, C.-Y., Bird, A.J., Winge, D.R. and Eide, D.J. (2007) Regulation of the yeast TSA1 peroxiredoxin by ZAP1 is an adaptive response to the oxidative stress of zinc deficiency. Journal of Biological Chemistry, 282, 2184–2195.

81. Wu, C.-Y., Roje, S., Sandoval, F.J., Bird, A.J., Winge, D.R. and Eide, D.J. (2009) Repression of sulfate assimilation is an adaptive response of yeast to the oxidative stress of zinc deficiency. Journal of Biological Chemistry, 284, 27544–27556.

82. Wu, C.-Y., Steffen, J. and Eide, D.J. (2009) Cytosolic superoxide dismutase (SOD1) is critical for tolerating the oxidative stress of zinc deficiency in yeast. PloS one, 4, e7061.

83. Rudolph, H.K., Antebi, A., Fink, G.R., Buckley, C.M., Dorman, T.E., LeVitre, J., Davidow, L.S., Mao, J.I. and Moir, D.T. (1989) The yeast secretory pathway is perturbed by mutations in PMR1, a member of a Ca2+ ATPase family. Cell, 58, 133–145.

84. Mandal, D., Woolf, T.B. and Rao, R. (2000) Manganese selectivity of pmr1, the yeast secretory pathway ion pump, is defined by residue gln783 in transmembrane segment 6. Residue Asp778 is essential for cation transport. The Journal of biological chemistry, 275, 23933–23938.

85. Zencir, S., Dilg, D., Shore, D. and Albert, B. (2022) Pitfalls in using phenanthroline to study the causal relationship between promoter nucleosome acetylation and transcription. Nat Commun, 13, 3726.

86. Grigull, J., Mnaimneh, S., Pootoolal, J., Robinson, M.D. and Hughes, T.R. (2004) Genome-wide analysis of mRNA stability using transcription inhibitors and microarrays reveals posttranscriptional control of ribosome biogenesis factors. Mol Cell Biol, 24, 5534–5547.

87. Sun, M., Schwalb, B., Pirkl, N., Maier, Kerstin C., Schenk, A., Failmezger, H., Tresch, A. and Cramer, P. (2013) Global Analysis of Eukaryotic mRNA Degradation Reveals Xrn1-Dependent Buffering of Transcript Levels. Molecular Cell, 52, 52–62.

88. D’Aurora, V., Stern, A.M. and Sigman, D.S. (1978) 1,10-Phenanthroline-cuprous ion complex, a potent inhibitor of DNA and RNA polymerases. Biochem Biophys Res Commun, 80, 1025–1032.

89. Reimann, C.W., Block, S. and Perloff, A. (1966) The Crystal and Molecular Structure of Dichloro (1, 10-phenanthroline) zinc. Inorganic Chemistry, 5, 1185–1189.

90. Niyogi, S.K., Feldman, R.P. and Hoffman, D.J. (1981) Selective effects of metal ions on RNA synthesis rates. Toxicology, 22, 9–21.

91. Lin, S.-J. and Culotta, V.C. (1996) Suppression of oxidative damage by Saccharomyces cerevisiae ATX2, which encodes a manganese-trafficking protein that localizes to Golgi-like vesicles. Molecular and Cellular Biology, 16, 6303–6312.

92. Portnoy, M.E., Liu, X.F. and Culotta, V.C. (2000) Saccharomyces cerevisiae expresses three functionally distinct homologues of the nramp family of metal transporters. Mol Cell Biol, 20, 7893–7902.

93. Jensen, L.T., Ajua-Alemanji, M. and Culotta, V.C. (2003) The Saccharomyces cerevisiae high affinity phosphate transporter encoded by PHO84 also functions in manganese homeostasis. Journal of Biological Chemistry, 278, 42036–42040.

94. Mei, Y., Jensen, L.T., Gardner, A.J. and Culotta, V.C. (2005) Manganese toxicity and Saccharomyces cerevisiae Mam3p, a member of the ACDP (ancient conserved domain protein) family. Biochemical Journal, 386, 479–487.

95. Reddi, A.R., Jensen, L.T., Naranuntarat, A., Rosenfeld, L., Leung, E., Shah, R. and Culotta, V.C. (2009) The overlapping roles of manganese and Cu/Zn SOD in oxidative stress protection. Free Radic Biol Med, 46, 154–162.

96. Walmacq, C., Kireeva, M.L., Irvin, J., Nedialkov, Y., Lubkowska, L., Malagon, F., Strathern, J.N. and Kashlev, M. (2009) Rpb9 subunit controls transcription fidelity by delaying NTP sequestration in RNA polymerase II. The Journal of biological chemistry, 284, 19601–19612.

97. Urban, J., Soulard, A., Huber, A., Lippman, S., Mukhopadhyay, D., Deloche, O., Wanke, V., Anrather, D., Ammerer, G. and Riezman, H. (2007) Sch9 is a major target of TORC1 in Saccharomyces cerevisiae. Molecular cell, 26, 663–674.

98. Pecci, L., Montefoschi, G., Musci, G. and Cavallini, D. (1997) Novel findings on the copper catalysed oxidation of cysteine. Amino Acids, 13, 355–367.

99. Prudent, M. and Girault, H.H. (2009) The role of copper in cysteine oxidation: study of intra- and inter-molecular reactions in mass spectrometry. Metallomics, 1, 157–165.

100. Fetherolf, M.M., Boyd, S.D., Taylor, A.B., Kim, H.J., Wohlschlegel, J.A., Blackburn, N.J., Hart, P.J., Winge, D.R. and Winkler, D.D. (2017) Copper-zinc superoxide dismutase is activated through a sulfenic acid intermediate at a copper ion entry site. The Journal of biological chemistry, 292, 12025–12040.

101. Sergio, S., Pirali, G., White, R. and Parenti, F. (1975) Lipiarmycin, a new antibiotic from Actinoplanes III. Mechanism of action. The Journal of antibiotics, 28, 543–549.

102. Talpaert, M., Campagnari, F. and Clerici, L. (1975) Lipiarmycin: an antibiotic inhibiting nucleic acid polymerases. Biochem Biophys Res Commun, 63, 328–334.

103. Sonenshein, A.L. and Alexander, H.B. (1979) Initiation of transcription in vitro is inhibited by lipiarmycin. Journal of molecular biology, 127, 55–72.

104. Srivastava, A., Talaue, M., Liu, S., Degen, D., Ebright, R.Y., Sineva, E., Chakraborty, A., Druzhinin, S.Y., Chatterjee, S., Mukhopadhyay, J. et al. (2011) New target for inhibition of bacterial RNA polymerase: ‘switch region’. Curr Opin Microbiol, 14, 532–543.

105. Chakraborty, A., Wang, D., Ebright, Y.W., Korlann, Y., Kortkhonjia, E., Kim, T., Chowdhury, S., Wigneshweraraj, S., Irschik, H., Jansen, R. et al. (2012) Opening and closing of the bacterial RNA polymerase clamp. Science, 337, 591–595.

106. Adams, C.C. and Gross, D.S. (1991) The yeast heat shock response is induced by conversion of cells to spheroplasts and by potent transcriptional inhibitors. Journal of bacteriology, 173, 7429–7435.

107. Nayak, D., Voss, M., Windgassen, T., Mooney, R.A. and Landick, R. (2013) Cys-pair reporters detect a constrained trigger loop in a paused RNA polymerase. Mol Cell, 50, 882–893.

108. Hein, P.P., Kolb, K.E., Windgassen, T., Bellecourt, M.J., Darst, S.A., Mooney, R.A. and Landick, R. (2014) RNA polymerase pausing and nascent-RNA structure formation are linked through clamp-domain movement. Nature structural & molecular biology, 21, 794–802.

109. Werner, F. (2012) A nexus for gene expression-molecular mechanisms of Spt5 and NusG in the three domains of life. J Mol Biol, 417, 13–27.

110. Benjamini, Y., Krieger, A.M. and Yekutieli, D. (2006) Adaptive linear step-up procedures that control the false discovery rate. Biometrika, 93, 491–507.

